# Design of bacterial DNT sensors based on computational models

**DOI:** 10.1101/2024.10.04.616532

**Authors:** Shir Bahiri Elitzur, Etai Shpigel, Itai Katzir, Uri Alon, Shimshon Belkin, Tamir Tuller

**Affiliations:** Department of Biomedical Engineering, Tel-Aviv University; Department of Plant and Environmental Sciences, Institute of Life Sciences, The Hebrew University of Jerusalem; Department of Molecular Cell Biology, Weizmann Institute of Science; Sagol School of Neuroscience, Tel-Aviv University

**Keywords:** Biosensors, computational biology, gene expression, landmines, explosives, synthetic biology, bioluminescence

## Abstract

Detecting explosive compounds such as 2,4,6-trinitrotoluene (TNT) and its volatile byproduct 2,4-dinitrotoluene (DNT) is paramount for public health and environmental safety. In this study, we present the successful application of diverse computational and data analysis models toward developing a bacterial biosensor engineered to detect DNT with high sensitivity and specificity. The *Escherichia coli-*based biosensor harbors a plasmid-based fusion of a gene promoter acting as the sensing element to a microbial bioluminescence gene cassette as the reporter. By analyzing endogenous and heterologous promoter data under conditions of DNT exposure, a total of 367 novel variants were generated. The biosensors engineered with these modifications demonstrated a remarkable amplification of up to 4-fold change in signal intensity upon exposure to 2,4-dinitrotoluene compared to non-modified biosensors, accompanied by a decrease in the detection threshold. Our analysis suggests that the sequence features with the highest contribution to biosensor performances are DNA folding patterns and nucleotide motifs associated with DNT sensing. These computational insights guided the rational design of the biosensor, leading to significantly improved DNT detection capabilities compared to the previous biosensor strain.

Our results demonstrate the effectiveness of integrating computational modeling with synthetic biology techniques to develop advanced biosensors tailored for environmental monitoring applications. A similar approach may be applied to a wide array of ecological, industrial, and medical sensing endeavors.

## Background

The detection of unexploded antipersonnel mines is a pressing global concern, impacting millions worldwide. Despite efforts by developed nations and humanitarian organizations to clear these mines, they continue to pose a significant threat, resulting in an estimated 15,000-20,000 casualties annually[1]. Beyond the humanitarian aspects, the presence of landmines disrupts various facets of life, ranging from agricultural activities to commercial endeavors. Despite ongoing clearance initiatives, the sheer magnitude of the task suggests that eliminating all current existing mines may require as long as 450-500 years at the current rate[2,3].

Presently, the predominant technologies for landmine detection mirror those employed during World War II, typically involving the use of metal detectors[4]. Not only do these methods yield high false positive rates, but they also fail to detect non-metallic munitions, which constitute most modern landmines. Moreover, the need for the presence of personnel on the actual minefield poses a significant risk of injury or death[5]. At present, there is no viable technology for the standoff detection of buried explosives.

In response to these challenges, alternative methods have been pursued, aiming to significantly reduce the false positive rate while maintaining a high probability of detection, thereby enhancing efficiency and minimizing the risk of harm. Such approaches encompass electromagnetic, acoustic, and explosive vapor sensing technologies, among others[6,7].^8^ Recently, the use of genetically engineered bacteria has been reported as explosive vapor’s sensors. This *E. coli*-based biosensor, activated by exposure to 2,4,6-trinitrotoluene (TNT) and 2,4-dinitrotoluene (DNT), employs a modified version of the *yqjF* gene promoter (C55)[8,9] as the sensing element, coupled with bioluminescent proteins encoded by a bacterial (*Aliivibrio fischeri*) *luxCDABEG* gene cassette[10] as the reporter element. However, despite these advancements, the performance of the sensor can be further improved.

A biosensor detects and records physiological or biochemical changes, relying on the specificity of its biological material. Key features include specificity, sensitivity, and real-time analysis. Microorganisms, particularly bacteria, are advantageous as biosensors due to their adaptability, ability to metabolize diverse compounds, and ease of genetic manipulation. Bacterial biosensor technology excels in tracking natural signaling pathways by controlling gene expression, allowing precise monitoring of cellular responses. This technology has been widely studied in the literature[11–21].

The aforementioned studies emphasize the effectiveness of bacterial biosensing, as evidenced by compelling proofs of concept. Specifically, the utilization of genetically engineered microbial biosensors presents a promising biological approach for the remote detection of landmines[22]. The concept revolves around the use of bacteria to generate an optical signal (reporter gene expression) in the presence of explosive vapors, enabling the detection of explosive devices from a safe distance[23,24]. Over the years, several strains of explosives-sensing bacteria have been reported, culminating in recent advancements such as the remote detection of buried antipersonnel landmines using an *Escherichia coli-*based biosensor[25–31].

Previous research has demonstrated how computational models can be used to analyze and solve complex problems in synthetic biology. For instance, one important aspect is predicting how molecules such as proteins and nucleic acids interact from a thermodynamic perspective[32]; kinetic models are also commonly used to control gene expression over time, allowing for the design of complex behaviors such as oscillations[33]. Another important area where such models can have an impact is the design of regulatory circuits, which are crucial for controlling cellular processes[34–36]. Computational models also help to predict changes in microbial metabolism to improve bioprocess productivity[37–40]. When we insert foreign genes into an organism, it is vital to construct stable genetic constructs that remain functional for extended periods[41]. Computational models also help us address other critical aspects of synthetic biology[42–44].

An alternative computational approach involves utilizing endogenous gene expression data and transcripts[45–48], due to their reliability. However, endogenous transcripts are influenced by various forms of evolutionary selection and possess diverse characteristics such as amino acid content and promoters, making it challenging to establish causality solely through their analysis. To address these challenges, an analysis of synthetic biology libraries may be employed. For instance, a study conducted to investigate how protein abundance is encoded in different transcript regions enabled researchers to identify new causal relationships between features in previously unexplored regions of transcripts and natural gene expression patterns[49]. Another study demonstrated how synthetic libraries can predict the fine-tuning of synthetic gene expression systems in yeast by incorporating various predictive intron features such as transcript folding and sequence motifs[50]. Researchers have also created synthetic promoter libraries to enhance the expression of desired proteins[51]. While numerous papers emphasize the effectiveness of utilizing synthetic biology libraries in research[52,53], to the best of our knowledge no such approach has been used in the field of biosensors.

The present study’s objective was to enhance the explosives’ detection capabilities of an existing microbial biosensor strain to improve its amenability to field requirements. To our knowledge, this study marks the inaugural utilization of sophisticated computational modeling of gene expression and gene design algorithms, with the combination of state-of-the-art synthetic biology methodologies, for the construction of a whole-cell biosensor[54]. While our primary focus lies in detecting buried landmines as a prototype system, the adaptable nature of this approach renders it applicable to a broad spectrum of environmental, industrial, and medical sensing scenarios.

## Results

### Optimization of the *yqjF* gene promoter

An optimization algorithm was employed to enhance the DNT sensing capabilities of the whole-cell biosensor. The procedure targeted the sensing element, the C55 version of the *yqjF E. coli* gene promotor[31,55], located upstream to the plasmid-borne *Aliivibrio fischeri* bioluminescence *luxCDABEG* gene cassette (Figure 1A). These genes encode 6 proteins, two of which (*luxAB*) code for the formation of the two subunits of the luciferase enzyme; the *luxCDE* gene products are responsible for substrate formation and recycling, and *luxG* for the supply of reducing power in the form of FMNH_2_. The computational optimization algorithm was comprised of several sequential steps, as detailed in the Method section and illustrated in Figure 1B. These steps involved analyzing both endogenous and heterologous promoter data under conditions of exposure and non-exposure to DNT. Subsequently, sequence motifs that were extracted distinguish between motifs in genes that were either overexpressed or under-expressed due to DNT exposure, utilizing a motif finding algorithm. The algorithm then computes the similarity of motifs between the two datasets, to assess the robustness of the DNT effect. It also examined these motifs’ positioning, to identify the optimal insertion site within the optimized promoter based on improvements in the position-specific scoring matrix (PSSM). Finally, the algorithm determines the final motif sequence by improving the PSSM score.

**Figure 1.**
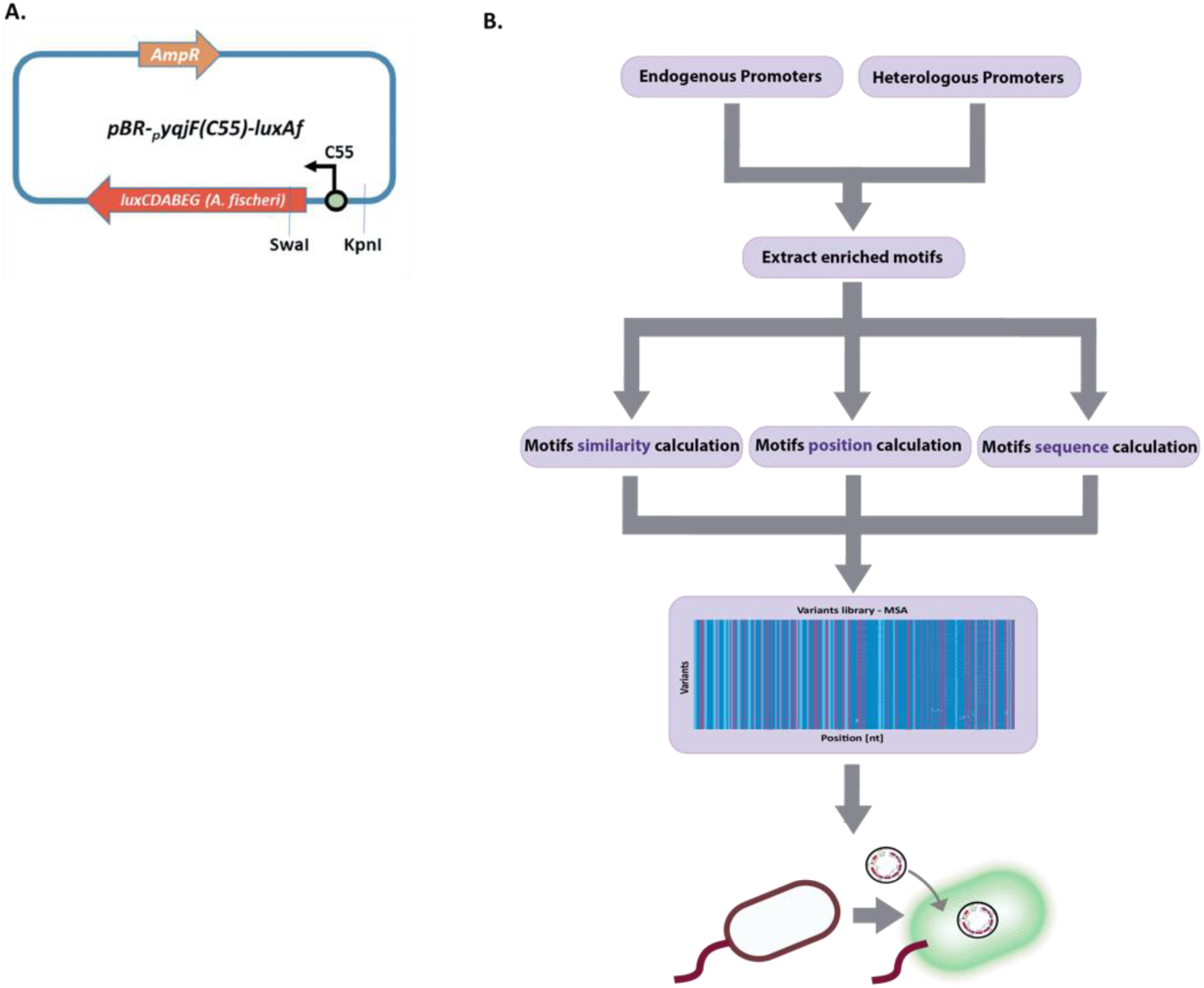
**A**. Scheme of the plasmid used in this study (pBR-C55-luxAf) with the KpnI and SwaI restriction sites. The yqjF (C55 version) gene promoter optimized during the research is marked. **B**. An illustration of the computational pipeline employed.

After this iterative process, a library comprising 397 variants of the *yqjF* gene promoter sequence was designed, out of which 367 plasmids were successfully synthesized and transformed into *E. coli (Supplementary Table S1)*.

The new biosensor strains thus generated were exposed to DNT (5 mg/L) for 10 hours (Figure 2A), and the maximum luminescence difference in the presence or absence of DNT was recorded (Figure 2B). Notably, 197 variants (over 53%) exhibited higher maximum luminescence values compared to the unmodified parental strain (illustrated by the purple line). Luminescence development by the top 5 performers in response to DNT exposure is depicted in Figure 2C; all five variants consistently outperformed the control throughout the experiment, in terms of both signal intensity and response time. These findings underscore the significant enhancement in the biosensor’s ability to detect explosive materials through the utilization of computational models to optimize gene expression.

**Figure 2.**
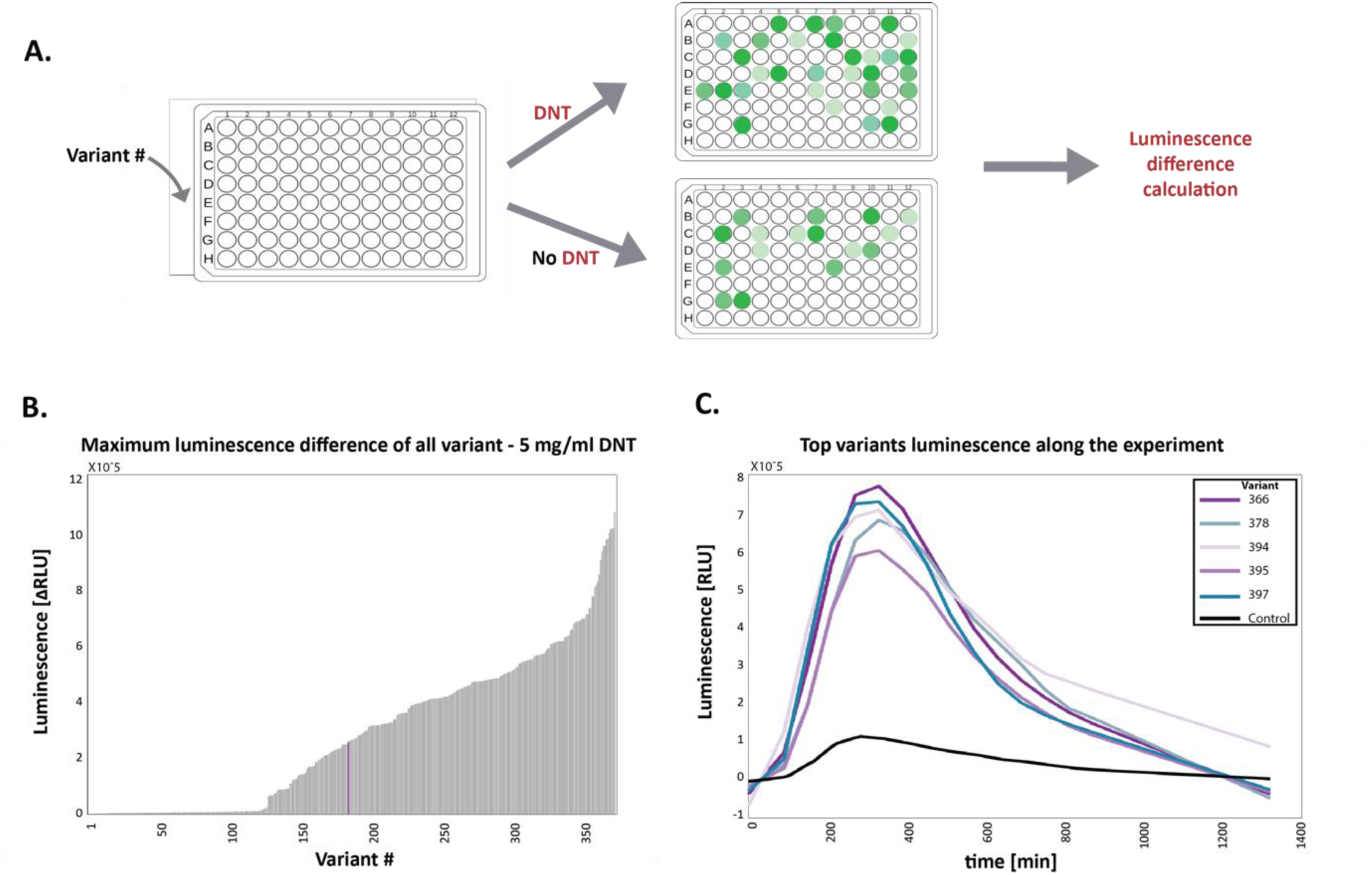
Library scan for response to DNT at 25°C. **A.** Experiment illustration. **B**. All variants are sorted by maximum luminescence difference net value. The purple line represents the unmodified control variant (average of three repeats). **C.** Luminescence development over time of the top 5 variants and the unmodified control variant (extrapolation of the experiment data to match all the time points of all experiments, more explanations in the Methods section). Luminescence data are provided in the plate reader’s arbitrary relative luminescence units (RLU).

### Evaluating the DNT detection threshold

To evaluate the effect of the *yqjF* promoter sequence modifications on DNT detection sensitivity, the top 5 variants were exposed to a series of DNT concentrations, along with the unmodified control. As depicted in Figure 3A, luminescence in the presence of all DNT concentrations was higher in the new variants; this was particularly apparent at the lower concentrations.

**Figure 3.**
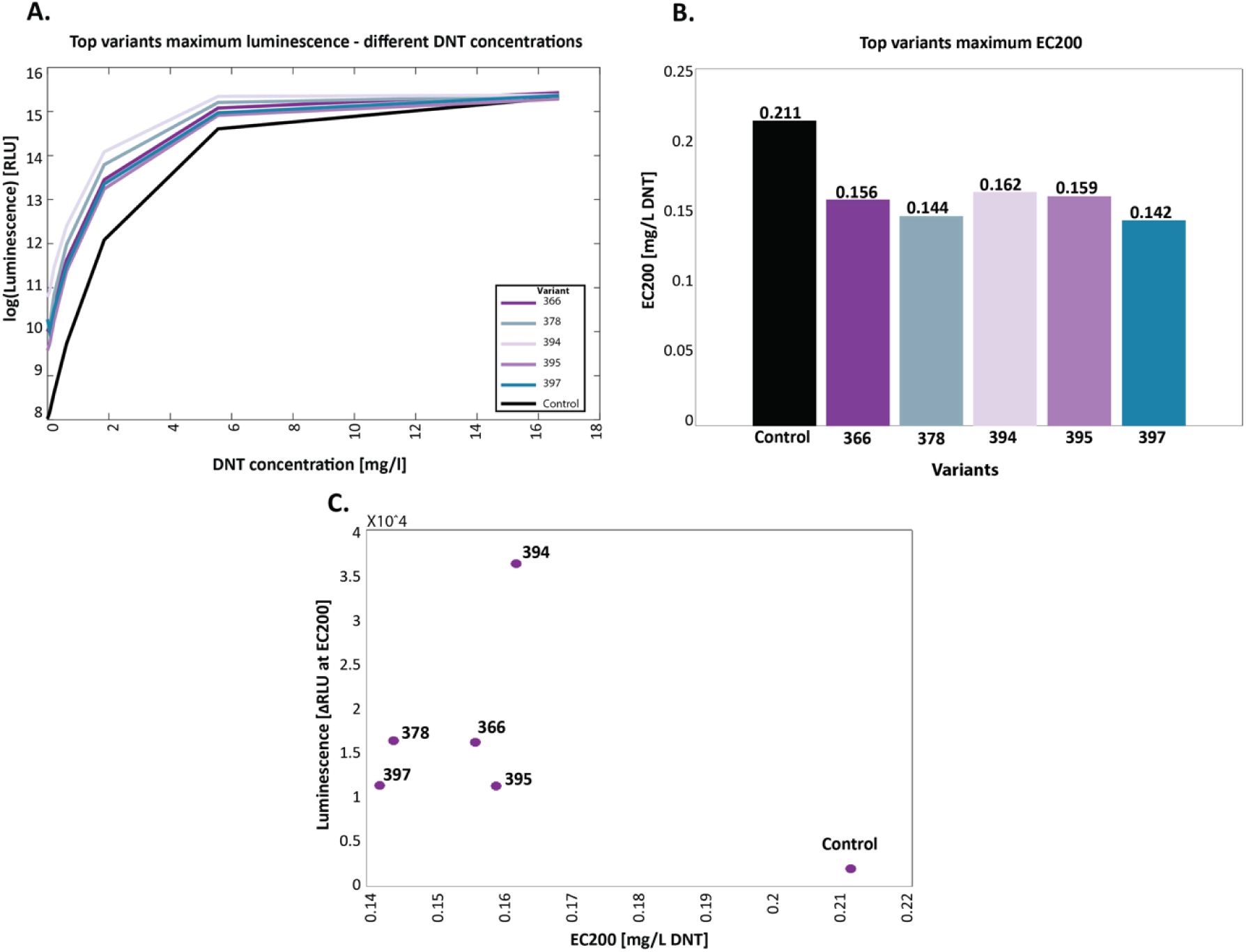
**A** Bioluminescent response to DNT of top-variant strains at 25°C. **A**. Maximum luminescence over a 5-hour exposure as a function of DNT concentration. **B.** Detection threshold, presented as the EC_200_ value (DNT concentration eliciting a 2-fold increase in bioluminescence compared to the uninduced control). **C.** Net signal intensity (ΔRLU, luminescence in the presence of DNT minus that in its absence) at a DNT concentration equal to EC_200_, plotted against the EC_200_ values.

To quantify these differences in sensitivity, the DNT detection threshold was estimated by calculating the EC_200_ value, denoting the DNT concentration eliciting a bioluminescent response that is 2-fold higher than the uninduced control. The EC_200_ values (Figure 3B) clearly demonstrate the enhanced detection sensitivity endowed by all 5 modified promoters. The improvements in both detection threshold and signal intensity are summarized in Figure 3C, in which luminescence intensity at the EC_200_ DNT concentration is plotted as a function of the EC_200_ value.

### Computational analysis of the variants

Three models were employed to analyze the predictive variables, as described in the Methods section: Linear regression, Lasso (Least Absolute Shrinkage and Selection Operator), and XGBoost (eXtreme Gradient Boosting). Figure 4A presents the cross-validation results, highlighting XGBoost as the model with the highest correlation in the validation set. Subsequent analyses were therefore based exclusively on this model.

**Figure 4.**
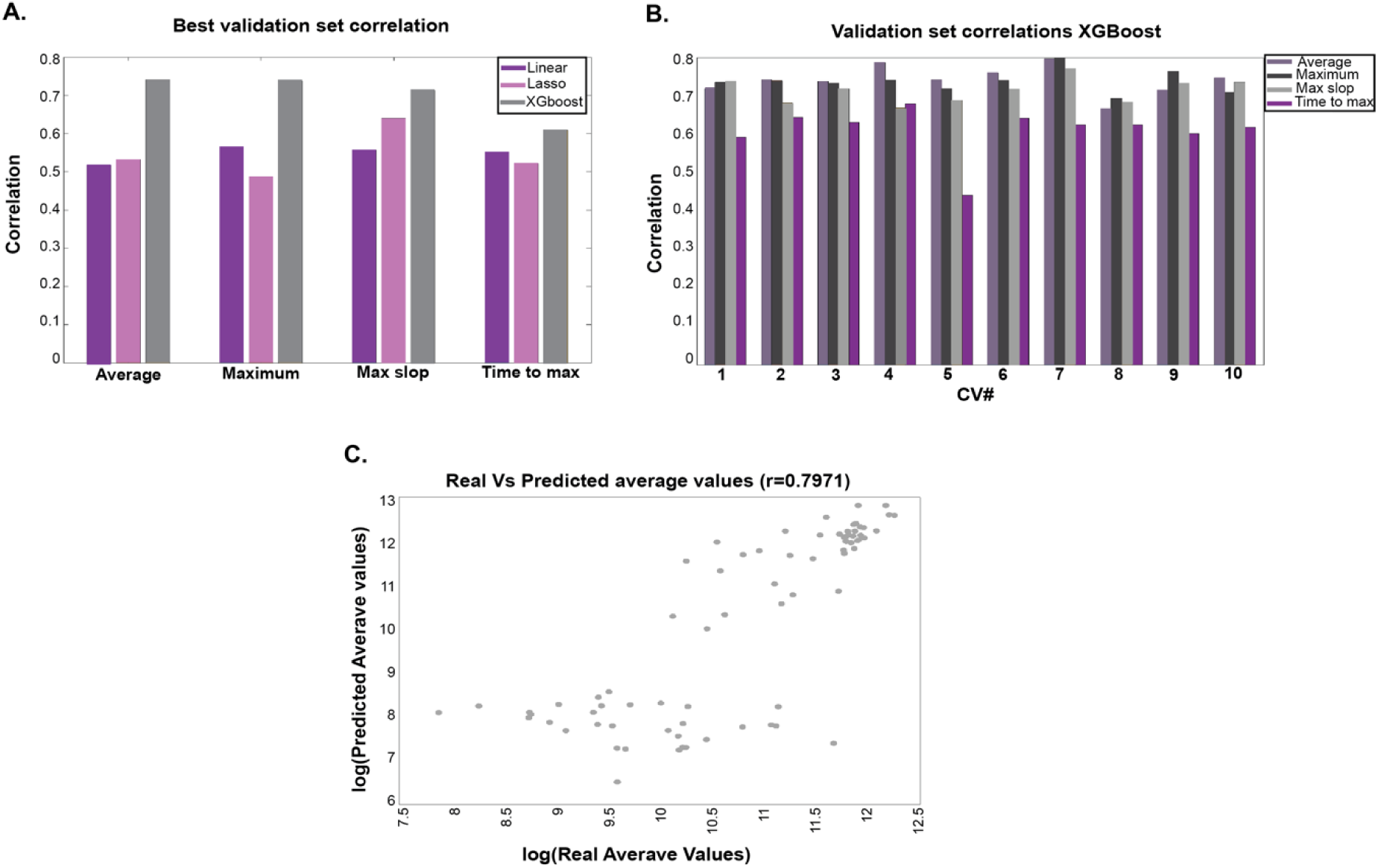
Prediction algorithm results. **A.** Best cross-validation, validation set correlation for all predicted variables. **B**. XGBoost validation set correlation for all cross-validation and predicted variables. C. Dot plot of the real and predicted variable “Average value”.

Figure 4B illustrates the correlation results for all cross-validation iterations across the different predictive variables, and Figure 4C displays the scatter plot for the variable “Average value”. Additional data are provided in the Supplementary Materials.

The results indicate that our algorithm can predict the main variables with high precision. Even when correlating the two repeats of the experiment, a correlation of 0.83 was achieved, demonstrating the robustness of our predictive model.

Given the strong correlation obtained, the next step was to analyze the features contributing to these results for each of the predictive variables. Figure 5 represents the “Average value” variable, and all other variables analyses can be seen in the Supplementary Figures S1, S2, and S3. First, we calculated the frequency of appearance of all features across all cross-validation iterations (Figure 5A, highlighting the top 10% of frequent features). Second, the XGBoost algorithm was utilized to rank the importance of features based on their F scores[56] (Figure 5B, presenting the top 30 features). Third, we employed SHAP (SHapley Additive exPlanations)[57] values to identify the most influential features and their direction of influence (Figure 5C and Figure 5D, showcasing the top 30 features). For more detailed information see the Method section.

**Figure 5.**
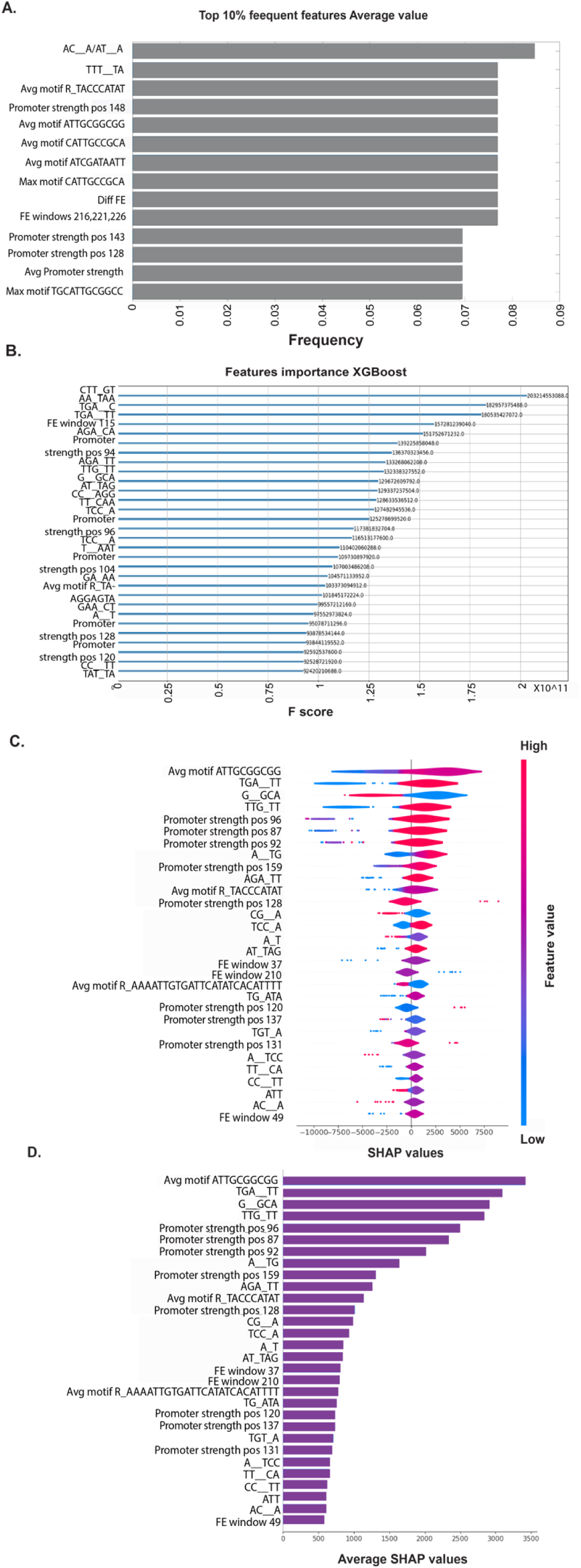
Most influential features predictor analysis. **A**. Top 10% of frequent features from all cross-validation sets. **B**. Top 30 features F score. C. Top 30 features SHAP values and direction. D. Top 30 features averaged SHAP values.

From this analysis, we observed that the most influential features were DNA folding in different windows, sequence motifs, and promoter strength. These findings are consistent with biological expectations: folding affects the ability to transcribe the coding sequence[58–60], while sequence motifs and promoter strength are crucial for the control and recruitment of transcription factors.[61] Notably, different techniques highlighted different important features, which is expected as each method considers different aspects of feature importance[62,63]. For example, when looking at our analysis of the frequency of features, it is affected by the cross-validation data division, while the SHAP values analysis focuses on the local and spatial explanations of the features, therefore different results can be obtained.

We have also correlated all important features identified by the various techniques to determine their consistency (Figure 6). We found that 50% of these features exhibited a high degree of correlation, indicating that they were robustly selected across different methods.

**Figure 6.**
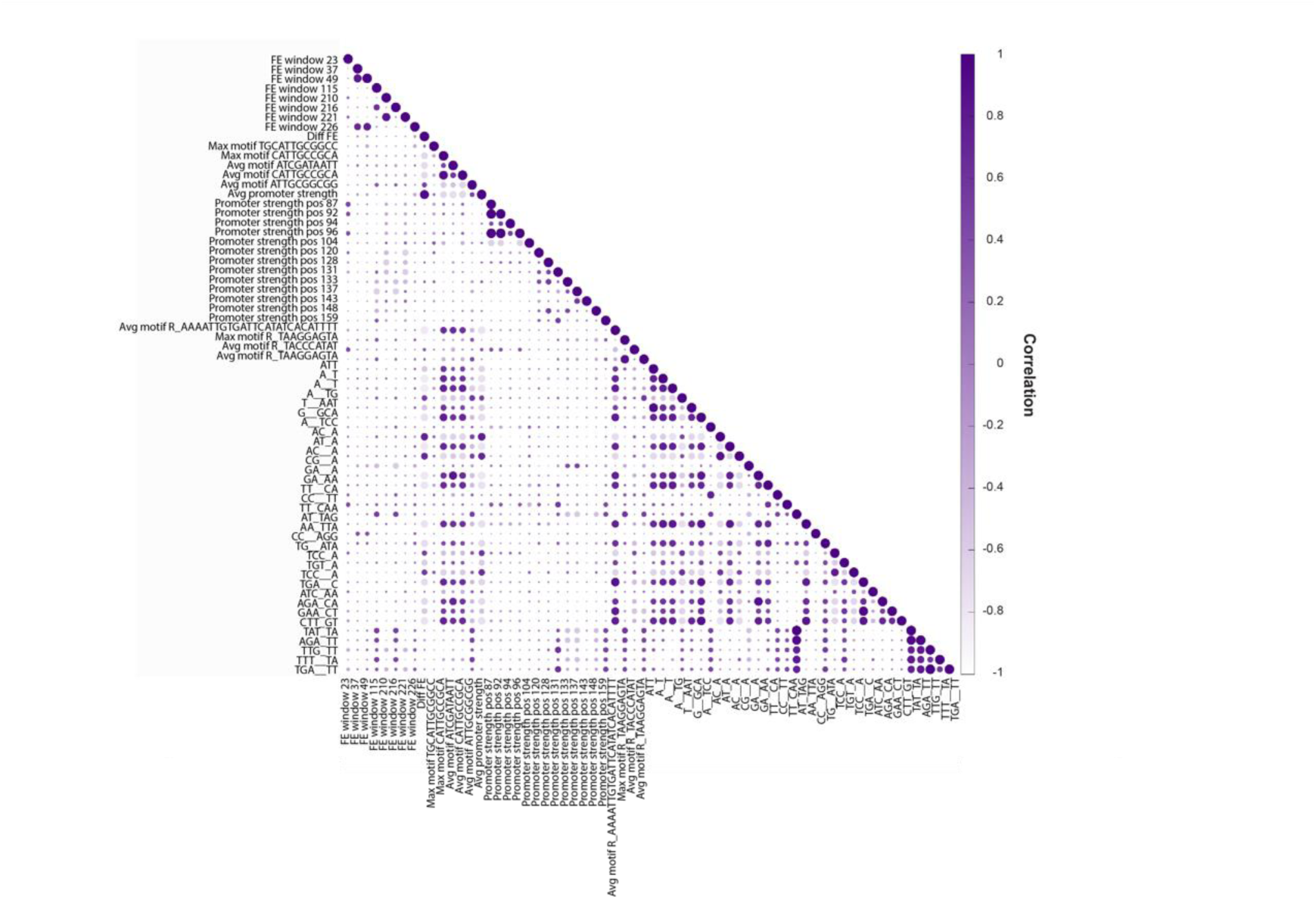
The Spearman correlation between each pair of the most important features was calculated and the correlation among the most important features is depicted, with the color and size of the circles indicating the strength of the correlation.

As shown in Figure 2B, the maximum luminescence difference differed significantly between the clones. Specifically, 53% of the variants exhibited higher values than the unmodified variant, while a substantial number of clones showed differences close to zero (indicating no observable change with and without DNT). This observation led us to create a classification algorithm to understand the reasons for this meaningful observation. The accuracy values obtained by employing the XGBoost and SVM (support vector machine) classifiers from all cross-validation iterations are shown in Figure 7A. The results were robust, with XGBoost achieving an average accuracy of 0.8027, outperforming SVM, which had an average accuracy of 0.6784. Consequently, subsequent analyses were focused on XGBoost.

**Figure 7.**
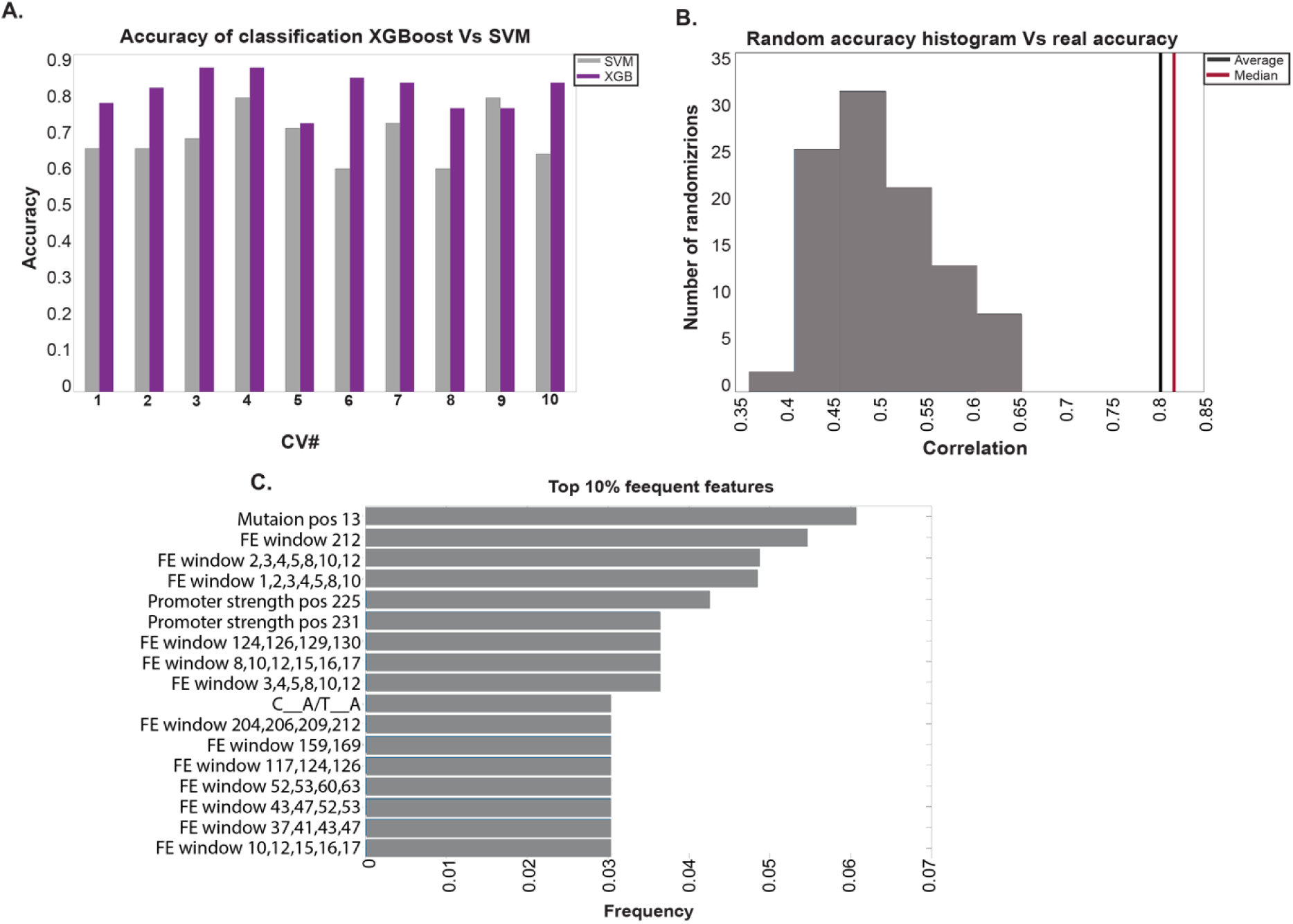
Classification algorithm results. **A**. XGBoost and SVM accuracy in all cross-validation sets. **B**. Random accuracy distribution and real accuracy average and median of all cross-validation sets. C. Top 10% frequent features of the classification, features labels are detailed in the Methods and Supplementary sections.

To validate the accuracy, we permuted the labels 100 times and reran the algorithm, calculating the accuracy for each randomization run. Figure 7B demonstrates that the classification accuracy with real labels was higher than all random accuracies.

We then examined the selected features by identifying the most frequent features across all cross-validation sets (Figure 7C, top 10% of frequent features). Notably, folding features had the most significant influence on the differences in maximum luminescence between variants.

## Discussion

In this study, we applied a computational-synthetic biology approach to improve the performance of a DNT biosensor, creating 367 variants that were evaluated against the best existing strain [53]. Prior strategies largely focused on molecular modifications, such as the directed evolution of the yqjF promoter[64] and its regulator YhaJ[65] as well as beneficial mutations introduced into the host bacterium[24]. Recently, additional manipulations of the YhaJ regulator[66] have further improved DNT detection capabilities. In the present communication, we highlight a different approach for sensor strain improvement, based on a computational approach that aims to identify motifs and features from sequences of variants or endogenous genes both exposed and unexposed to an explosive, and to infer which specific subsequences are affected and the reasons behind their influence. The result was significant, with 53% of the new variants exceeding the performance of the original strain, demonstrating the value of this approach.

The computational tools enabled the identification of crucial features affecting biosensor performance. DNA folding patterns emerged as critical to regulatory element accessibility, potentially allowing transcription factors to bind more efficiently. Additionally, specific nucleotide motifs were found to influence biosensor sensitivity by functioning as key recognition sites for transcription factors. These findings point to the broader significance of integrating computational techniques with synthetic biology to improve biosensor design.

Furthermore, this study opens the door to a deeper exploration of the metabolic pathway involved in DNT sensing. While both DNT and TNT activate the yqjF promoter, the data suggest that their degradation product, trihydroxytoluene[67], is the more likely inducer. This metabolic interaction, combined with the role of YhaJ in regulating yqjF activation[68], offers a compelling opportunity for further optimization of the biosensor[66], focusing on enhancing the biosensor’s sensitivity and performance by targeting these regulatory mechanisms.

## Conclusion

In conclusion, this work demonstrates the significant potential of computational approaches for the optimization of biosensors. By analyzing the structural and sequence motifs within the new variants, we were able to identify key features such as DNA folding patterns and nucleotide motifs that significantly enhanced biosensor performance. Importantly, our findings show that computationally designed biosensor variants can surpass traditional molecular modification strategies, with over half of the new variants outperforming the original strain. This result underscores the importance of integrating computational models to predict and identify regulatory elements that influence biosensor function.

The broader applicability of this methodology is evident, as the principles outlined here can be extended to a variety of biosensing applications, ranging from environmental monitoring to industrial and medical diagnostics. The ability to fine-tune sensor sensitivity and signal strength through computational means offers a scalable approach to biosensor design, providing a path for more adaptable and efficient systems. Additionally, the insights gained into the underlying metabolic pathway responsible for DNT sensing, particularly the potential role of trihydroxytoluene as a key inducer, highlight the need for further research into optimizing these metabolic interactions.

Future directions should aim at refining biosensor performance by enhancing the interactions between the sensor and the YhaJ transcription factor. This, combined with a deeper understanding of the metabolic processes involved in DNT detection, will likely lead to even more robust and sensitive biosensors. Overall, this study provides a solid foundation for future innovations in biosensor technology and reinforces the value of combining synthetic biology with computational models to address complex biological sensing challenges.

## Methods

### Chemicals and reagents

DNT (2,4-dinitrotoluene) was purchased from Sigma-Aldrich (cat. 101397). A working stock of 10 g/L in ethanol was prepared and kept at room temperature.

### Plasmids and strains

The previously described[53] plasmid pBR-C55-luxAf, harboring the *Aliivibrio fischeri luxCDABEG* gene cassette downstream of the *yqjF* gene promoter (version C55) served as the chassis for testing the different *yqjF* variants^69^ (Figure 1A). *E. coli* strain BW25113[71] served as the host in all experiments.

### The analyzed data libraries

To enhance the biosensor’s detection capability and sensitivity, two datasets were used: the endogenous genes dataset and the heterologous genes dataset. The goal was to address the limitations of each—specifically, resolving causality issues in the endogenous dataset and improving specificity in the heterologous dataset

### Endogenous library data

We utilized RNA-seq data from *E. coli* genes, both with and without exposure to DNT (5 mg/L)[72], to identify endogenous genes exhibiting either over-expression or under-expression upon DNT exposure. For each gene, we quantified the expression disparity and selected the promoters (each with a length of 400) corresponding to the over-expressed genes and a subset representing the under-expressed genes (10%). This dataset is denoted as Dataset A.

### Synthetic library data

The synthetic library, detailed in Table S2, is based on the *yqjF* gene promoter fused with GFP and includes single mutations across 147 sites, as well as various mutation pairs, triplets, and higher-order combinations, totaling approximately 100,000 sequences. To assess promoter activity, flow-seq method[73] was employed and sorted the library into 16 logarithmic bins based on GFP fluorescence using Fluorescence-Activated Cell Sorting (FACS), as shown in Figure S4. This sorting was performed twice, once with DNT (200 μg/ml) and once without.

To identify motifs potentially influencing expression in response to DNT exposure, we categorized the promoters into over-expressed and under-expressed groups, each comprising the top 10% based on the weighted average fluorescence of individual promoters:

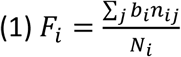

Where *b_i_* is the average fluorescence in bin i, nij is the number of appearances of variant j in bin i, and *N_i_* is the total count of variant i.

We refer to this data set as data set B.

### Sensing element (C55 promoter) optimization

To optimize the whole-cell biosensor and improve its DNT sensing capability, we focused on enhancing the C55 promoter within the plasmid, based on the *yqjF* gene promoter, as shown in Figure 1A. The optimization was tailored for each dataset and involved the following steps:

### Step 1 – Sequence Motif Extraction

We used a differential enrichment algorithm to discover significant motifs by comparing target and reference sets. Searches were conducted with the following configurations:

1. Over-expressed promoters as the target set and all promoters as the reference.
2. Over-expressed promoters as the target set and under-expressed promoters as the reference.
3. Under-expressed promoters as the target set and all promoters as the reference.
4. Under-expressed promoters as the target set and over-expressed promoters as the reference.

This process was applied separately to Dataset A and Dataset B, resulting in 159 significant motifs for Dataset A and 596 for Dataset B.

### Step 2 – Find similar motifs

To streamline the motif pool and ascertain consistency across motifs derived from both datasets (A and B), we aimed to identify similarities between motifs. Each motif was represented by a position-specific scoring matrix (PSSM). To facilitate comparison between motifs, a distance metric was employed:

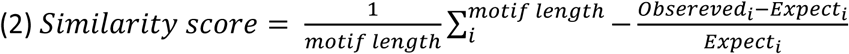

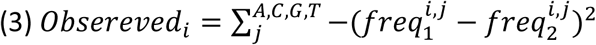

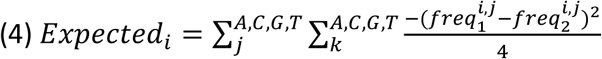

To compare PSSMs, we standardized the number of rows and columns by applying equal padding (0.25) to motifs of varying lengths. Padding was uniformly added along the edges in all configurations, as shown in Figure 8A, with the final score being the maximum similarity score.

**Figure 8.**
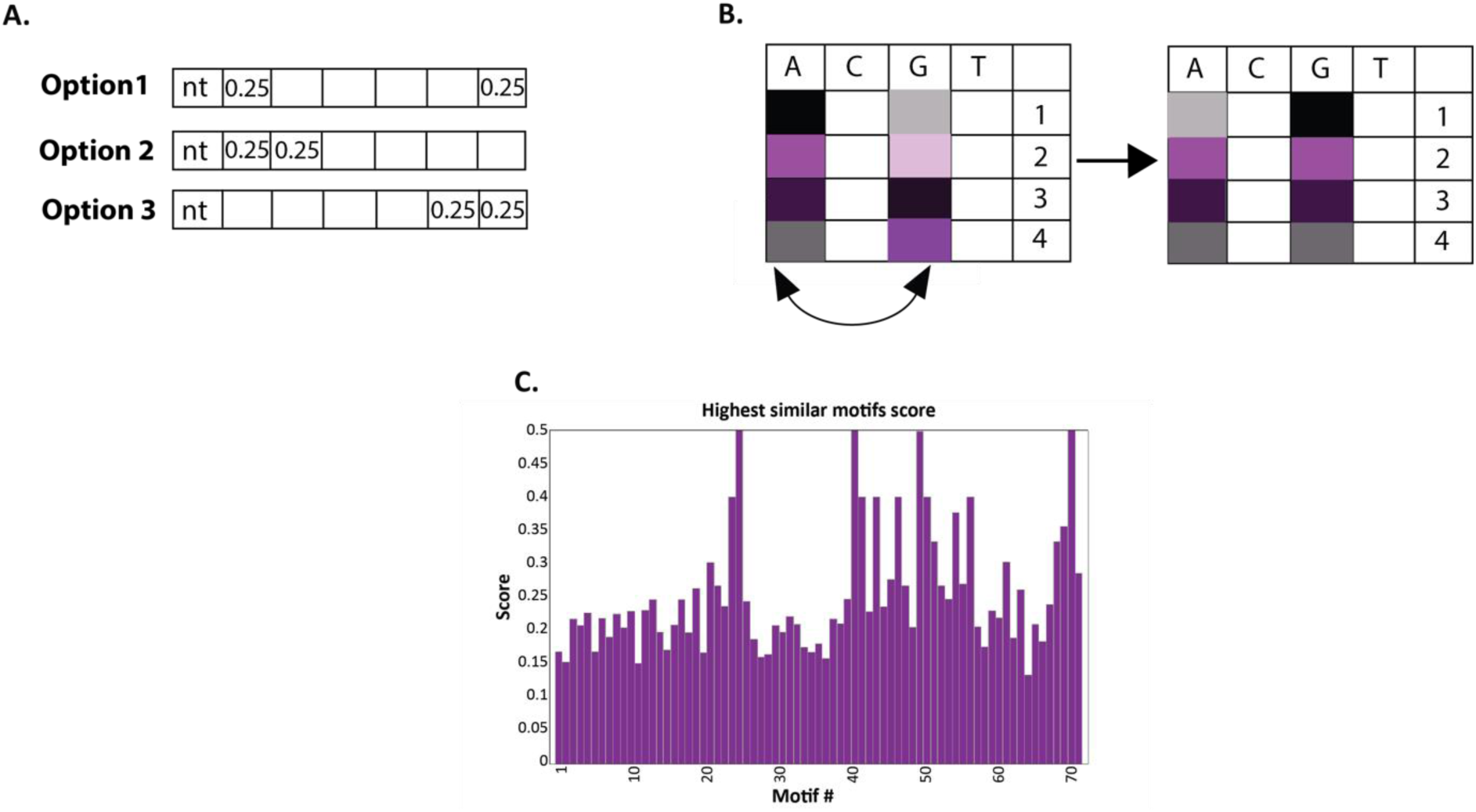
**A**. Motifs padding options. To equal the motif’s dimension, we padded the edges of the relevant motifs with equal probabilities for each nt. An example in the illustration is padding a motif with four letters to a motif with six. **B**. Permutation illustration. We permuted the nt distribution of the motifs PSSM to create a null model to find significant similarity scores (Method section). **C**. We calculated a similarity score between each pair of motifs (data set A and data set B). Only significant motifs with the highest similarity score were chosen. The bar graph shows the highest similarity score of the significant motifs.

To set a meaningful threshold, we used an empirical P-value test. We permuted the rows (nt) of the PSSMs 100 times and computed random similarity scores, as illustrated in Figure 8B. A score was considered significant if it was ≤ 0.05.

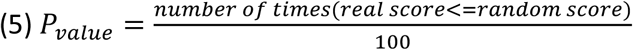

For each motif in Dataset A, we compared it with all motifs in Dataset B and selected the one with the highest similarity score. Only motifs with significant scores were retained, as shown in Figures 8A and 8B, with the highest similarity scores illustrated in Figure 8C. This process identified 73 motifs with significant similarity and relevance.

### Step 3 – Find significant motif positions

The discovered motif is linked to target and reference sequences and a PSSM matrix. To find the optimal insertion site within the C55 promoter, we maximize or minimize the PSSM score based on whether the motif is from over- or under-expressed promoters.

The process for identifying the most influential insertion site involves:

1. Selecting a motif.
2. Choosing a relevant promoter for the motif.
3. Calculate the PSSM score at all positions within the promoter using a sliding window.
4. Generating 100 permutations of the promoter by shuffling nucleotides while preserving nucleotide distribution and GC content.
5. Calculate the PSSM score for the motif at all positions in each permuted promoter.
6. Determining the empirical p-value for each position to identify significant sites.

This process is repeated for all motifs and their respective promoters, as shown in Figure 9A.

**Figure 9.**
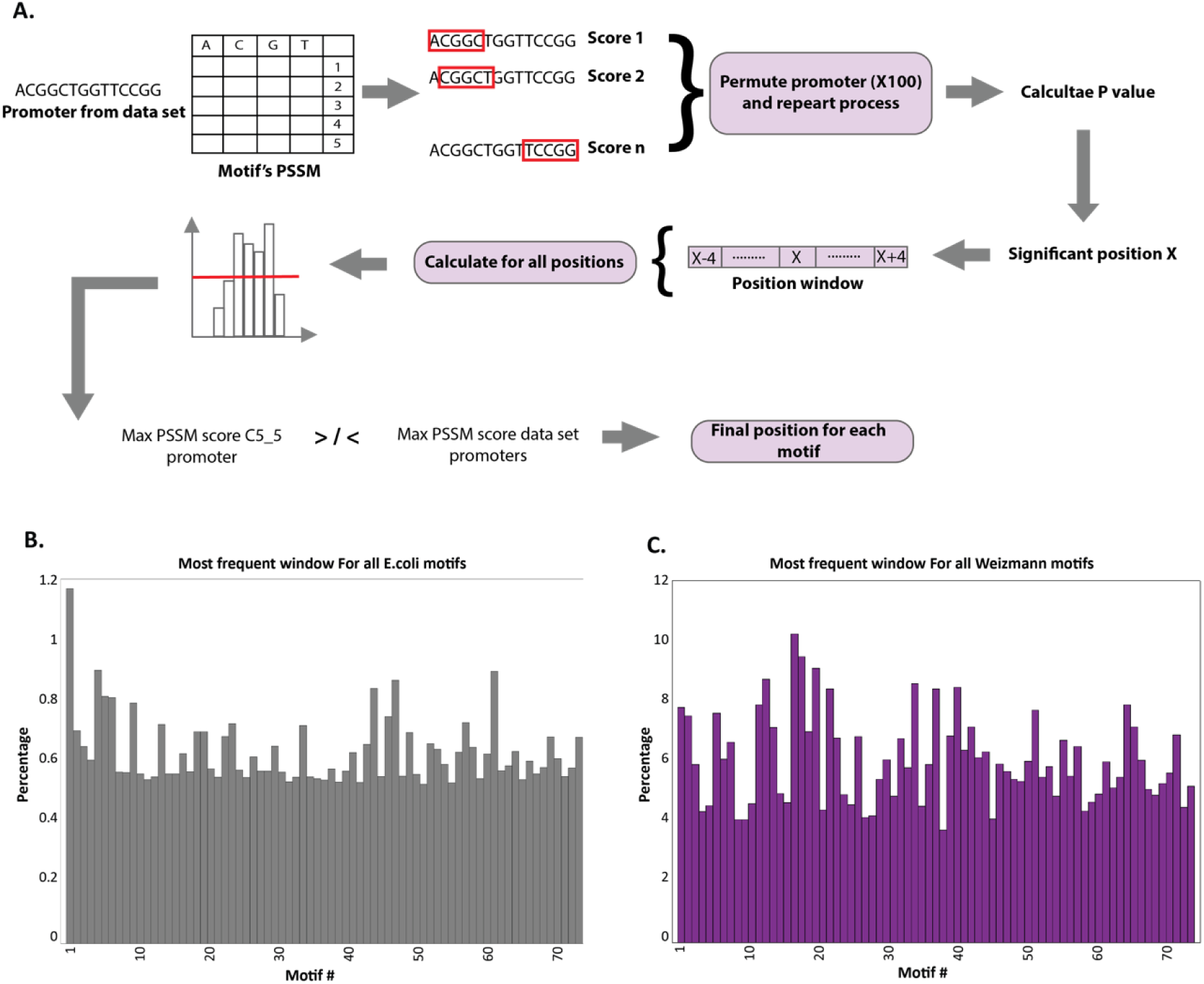
**A**. Illustration of the algorithm that finds significant positions of the motifs. **B**. Most frequent positions window for all motifs (Methods section) data set A. **C**. Most frequent positions window for all motifs (Methods section) data set B.

It is noteworthy that each motif may have multiple significant positions across different promoters. Thus, our objective was to identify the most prevalent position for each motif. To achieve this, the following additional steps were undertaken:

1. Calculate the frequency of occurrence (x_i) for each significant **position** across all relevant promoters.
2. Expand each **position** into a **window** spanning ±4 nucleotides to provide greater flexibility.

Step 3: Calculate the sum of appearances for each **window** using Equation 6:

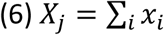

Step 4: Compute the percentage of appearances for each **window** using Equation 7:

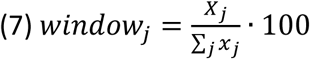

Step 5: Select the **window** with the maximum percentage of appearances for each motif (max(*window_j_*))

Motifs were selected if they appeared in windows with a percentage greater than 6.5%, ensuring a sufficient number of motifs. Due to lower percentages in Dataset A, motifs from this dataset were matched with similar motifs from Dataset B (see Figures 9B and 9C).

For motifs from both datasets, significant positions were those that were identical or within ±20 nt.

The final step was to determine the optimal insertion position for each motif by calculating and comparing PSSM scores at relevant positions within the C55 promoter and the corresponding promoters from both datasets, as outlined in Equations 8 and 9. The goal was to insert motifs from over-expressed promoters and remove those from under-expressed ones. Promoter lengths varied, so positions were aligned to the START codon.

For motifs from over-expressed promoters, the selected positions had to meet the following conditions:

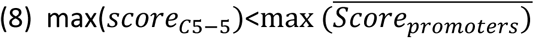

Where *score*_*C*5−5_ is a vector of PSSM’s scores of a motif in all its significant position in the C55 promoter, 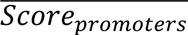 is the average score of PSSM’s scores of a motif in all its significant position and all relevant promoters.

For motifs from under-expressed promoters, the selected positions met the following conditions:

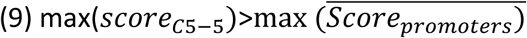

Where *score*;_*c*5−5_ is a vector of PSSM’s scores of a motif in all its significant position in the C55 promoter, 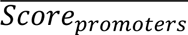 is the average score of PSSM’s scores of a motif in all its significant position and all relevant promoters.

After this process, we identified 13 motifs from Dataset A and 15 motifs from Dataset B.

### Step 4 – Find the final motif’s sequence

Since determining the consensus sequence of a motif from a PSSM matrix is not straightforward, we developed an algorithm to maximize the PSSM score of a motif at its designated position. The goal is to optimize the motif’s impact by maximizing its score, as outlined in Figure 10A.

**Figure 10.**
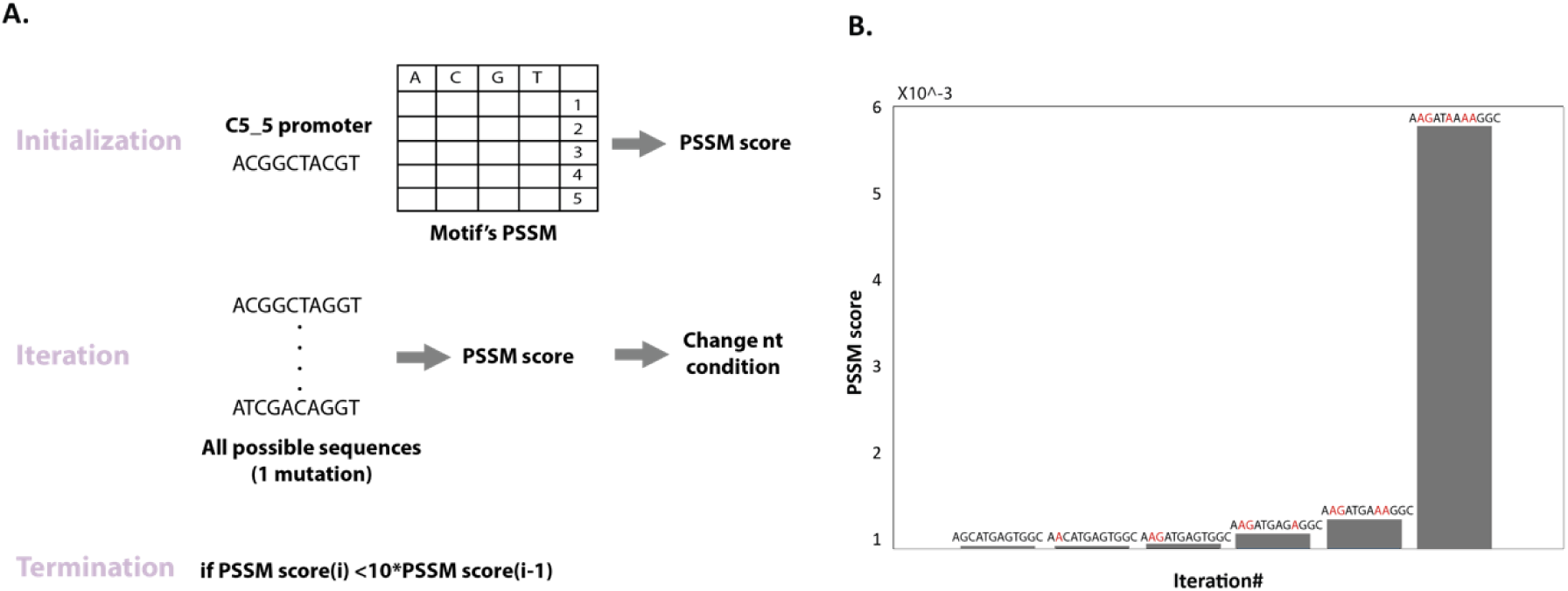
**A**. Illustration of the algorithm that finds motifs sequence (Method section). **B**. An example run of the algorithm. The graph shows the improvement of the PSSM score and the change in the nt sequence on each iteration.

Algorithm Steps:

1. Initialization: Compute the initial PSSM score of the motif at its position in the promoter.
2. Iteration:

- Generate a pool of sequences with single mutations.
- For each position within the motif, substitute the nucleotide with each of the other three possible nucleotides.
- Calculate the PSSM score for each mutated sequence.
- Select the sequence with the highest score change compared to the previous iteration.
- Repeat the process without altering the position changed in the last iteration.
3. Termination: The algorithm stops when no further improvements are observed, and the final sequence is chosen as the optimal motif sequence.

Figure 10B illustrates an iteration of the algorithm, showing score improvements and nucleotide changes.

### Step 5 – Insert motif into the promoter

This phase involves constructing a library of variants. Initially, positions that are constrained by factors such as restriction sites are identified and marked, precluding any alterations. Subsequently, each motif is individually inserted into the plasmid. Finally, motifs from both datasets are concurrently incorporated into the same variant using a Cartesian product approach, with the conditioning that they do not overlap.

#### Construction of the *yqjF* variant library

All variant *yqjF* promoter sequences were synthesized and cloned by Twist Bioscience (San Francisco, CA, USA) and were obtained pre-cloned, at KpnI/SwaI sites (Figure 3A), in a pBR-X-luxAf backbone (*Aliivibrio fischeri luxCDABEG*). Overall, 397 clones were ordered, 38 of which failed Twist Bioscience QC before synthesis. Hence, 367 clones were successfully synthesized and tested for their reaction to DNT.

#### Bioluminescence assays

Each clone was transformed separately into *E. coli* DH5α and grown overnight (37°C, 200 rpm rotary shaking), in 96-well plates, each well containing 150 µl lysogeny broth (LB) with 100 µg/ml ampicillin. Overnight culture aliquots were transferred into individual wells of duplicate 96-well plates, each well containing 50 µl of the same medium. Following 3 h growth under the same conditions, 50 µl of DNT dissolved in LB (final concentration 5 mg/L) was added to one plate and the same volume of DNT-free LB was added to the other. Both plates were incubated at 25°C for 10 hours, in the course of which bioluminescence and optical density at 600 nm were measured every hour using a Microlab Star robot (HAMILTON, Switzerland) and a Synergy HT, BioTek plate reader. The following values were recorded for each clone: maximum luminescence intensity, net luminescence (after subtraction of the control luminescence), and response ratio (luminescence divided by the control luminescence). The five best-performing clones were selected based on the following criteria: having both the highest net luminescence and the highest ratio along with faster signal development.

The response of the selected clones to DNT was analyzed by exposing them to a DNT concentration series and measuring the ensuing luminescence as a function of time.

grown overnight at 37°C with vigorous agitation in LB supplemented as before. The culture was diluted x1/100 in LB without antibiotics and regrown under the same conditions to an OD_600_ = 0.3. Bacterial aliquots (50 µl) were transferred to an opaque white 96-well plate (Greiner Bio-One), already containing 50 µl of DNT at various concentrations in 4% ethanol (2% final concentration). Bioluminescence was measured every 15 minutes at ambient temperature, using a microplate reader (either a TECAN Infinite® 200 PRO, Männedorf, Switzerland, Synergy HT, BioTek or a Wallac VICTOR2, Turku, Finland). All experiments were conducted with internal duplicates, were repeated at least three times and pBR-C55-luxAf served as the control strain.

Luminescence data are presented either as the instrument’s raw arbitrary relative light units (RLU) or as the net signal in the presence minus the absence of the inducer (ΔRLU). The detection sensitivity of the biosensors to DNT is presented as the EC_200_ value, representing the DNT concentration at which luminescence in the presence of DNT is twice that in its absence[74,75].

#### Computational analysis of the library

To analyze the variants’ behavior, we compared luminescence data from different plates, requiring interpolation and extrapolation to standardize measurements across consistent time points. As shown in Figure 2B, the variants displayed varied behaviors, prompting an investigation into their sequences to understand the luminescence results.

### Extracted features

We extracted 6,711 sequence-related features from the promoters of the variants, including the following categories (can be seen in Table S3):

1. Folding Energy: The folding energy of the variants’ promoters was calculated using the MATLAB function “rnafold”[76].
  1. Folding in Windows: Calculation of the folding energy for every 40 nucleotides with a sliding window of 1 (the window’s position is determined by the first nucleotide in the window).
  2. Average Folding: The average folding value across all windows.
  3. Total folding: the folding energy of the entire variant.
2. Mutations: Each promoter was compared to the control promoter to identify mutations.
  - Existence of Mutation: A binary feature indicating the presence (1) or absence (0) of a mutation at a specific position.
  - Amount of Mutation: The total number of mutations in a variant.
3. Chimera ARS (Average Repetitive Substring) Index[77]: This index is based on the propensity of coding regions to include long substrings that appear in other coding sequences, assumed to be regulatory regions. We calculated the average repetitive substring in our variants using a reference set (the top 20% of over-expressed *E. coli* promoters).
4. Sequence Motifs:
  1. PSSM Scores[78]: Maximum Position-Specific Scoring Matrix (PSSM) scores of the inserted motifs.
  2. New Motif Extraction: Extraction of new motifs as described in the Sequence Motifs Extraction section.
  3. Motifs from SwissRegulon[79]: Downloaded from the SwissRegulon *E. coli* database, which contains genome-wide annotations of regulatory motifs, promoters, and transcription factor binding sites (TFBSs). This set includes 37 motifs.
5. Promoter Strength[80]: Calculated using an energy matrix [KBT] from Brewster et al. The matrix covers base pairs [-41:-1], where 1 denotes the transcription start site. Each row corresponds to a nucleotide, and each column corresponds to a position.
  1. Strength by Position: Promoter strength is calculated for specific positions.
  2. Average Promoter Strength: Overall promoter strength across the entire sequence.
6. Nucleotide Count Features: Based on nucleotide counts from Muhammod et al[81]., as detailed in Table S3 in the Supplementary section.
7. Ribosome binding site strength: calculated by a high-resolution computational model that predicts rRNA-mRNA interactions’ strength in a sliding window with the size of six nt along the promoter[82].

This detailed extraction of sequence-related features enabled a comprehensive analysis of the variants’ promoters, facilitating the understanding of their luminescence behavior.

To streamline our feature set:

1. Folding in windows: we calculated the Spearman correlation between folding features and retained only one representative feature for those with a correlation above 0.99.
2. Mutations: We averaged mutation profiles at similar positions to reduce redundancy.

These steps improved the efficiency of our analyses.

### Prediction variables

We used K-means clustering with correlation as the distance metric to group variants based on luminescence differences over time. After testing various K values, we selected K = 2 based on cluster evaluation metrics (Davies–Bouldin index[83] and Silhouette[84] score). As can be seen in Figure 11, for K=2 the average luminescence differences pattern is different, and from that we chose to predict four variables: maximum luminescence difference, average luminescence difference, maximum slope value, and time to maximum luminescence difference.

**Figure 11.**
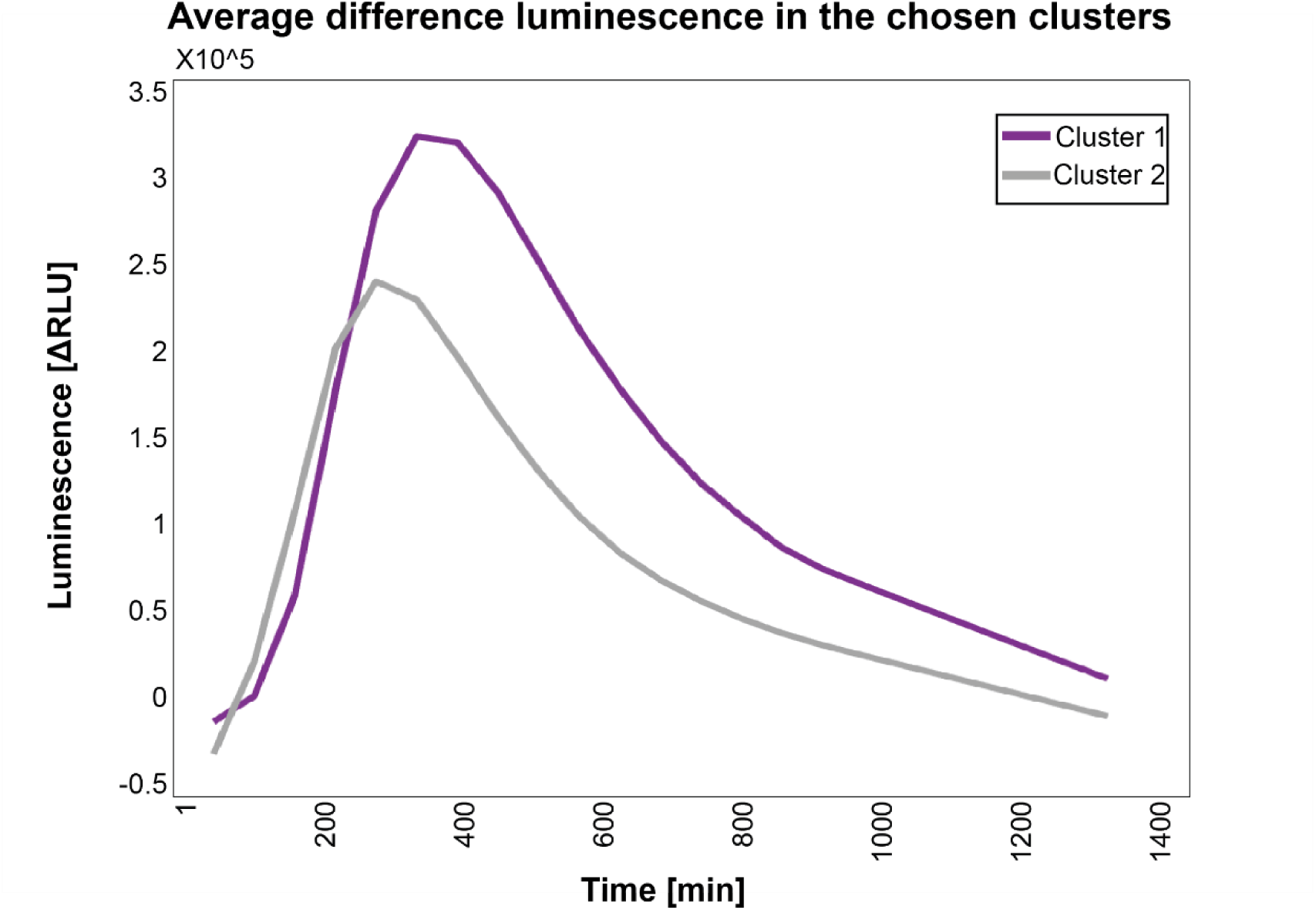
All luminescence difference values of the variants in each cluster were averaged to examine different trends in the groups.

### Predictor Algorithm

To predict the selected variables, we employed three regression models:

1. Linear Regression:

- The data was split into training (60%), testing (20%), and validation (20%) sets, with each cross-validation iteration using a different random split.
- Features were added using forward selection to maximize the correlation on the test set.
- The process was repeated for 10 cross-validation iterations.
2. Lasso Regression[85]:

- The data was divided into training (80%) and validation (20%) sets.
- The process was repeated for 10 cross-validation iterations.
- The “lassocv”[86] function in Python was used to optimize the Lasso algorithm’s parameters, iteratively fitting along a regularization path.
3. XGBoost (eXtreme Gradient Boosting[87]):

- The data was split into training (60%), testing (20%), and validation (20%) sets, with each cross-validation iteration using a different random split.
- Features were added using forward selection to maximize the correlation on the test set.
- The process was repeated for 10 cross-validation iterations.
- Hyperparameters were optimized using Optuna[88], which automates the search for optimal configurations through Bayesian optimization. Each trial used a different set of hyperparameters, evaluating the R-squared score on the validation set. The best hyperparameter set for our data and model was selected after adjusting the range and re-running Optuna (details provided in the supplementary section Figure S5).

This approach ensured that the most effective predictive model and feature set were selected, enhancing the accuracy and robustness of our predictions.

### Classification algorithm

As can be seen in Figure 2B there is a division between the maximum luminesces difference of the variants. To examine the difference, we classified the variant into 3 groups according to two thresholds (relative to the control variant, while having balanced data in each label group). We used two classification models:

1. SVM (support vector machine)[89]:

- Radial Basis Function (RBF) kernel
- The data was divided into training (80%) and validation (20%) sets with each cross-validation iteration using a different random split.
- Features were added using forward selection to maximize the accuracy of the test set
2. XGBoost:

- The data was split into training (80%), and testing (20%) sets, with each cross-validation iteration using a different random split.
- Features were added using forward selection to maximize the accuracy of the test set.
- The process was repeated for 10 cross-validation iterations.
- Hyperparameters were optimized using Optuna[88], which automates the search for optimal configurations through Bayesian optimization. Each trial used a different set of hyperparameters, evaluating the R-squared score on the validation set. The best hyperparameter set for our data and model was selected after adjusting the range and re-running Optuna.

To validate the classification results, we created 100 randomized versions of the labels (permuted the labels) and ran our classification models.

### The selected features

To identify the most meaningful features, we utilized and combined several methods:

1. Frequency of Features in Cross-Validation: We aggregated all selected features from the cross-validation runs and calculated their frequency, focusing on the top 10% most frequent features. It is important to note that we also reduced the number of features following the application of our algorithms:

- Folding Energy: We combined windows that are within 10 units of each other.
- Sequence Features: We merged features that have only a single mutation difference.
2. XGBoost Feature Importance: We ranked the importance of features using the XGBoost algorithm, which evaluates feature importance based on the F score[56]. The F score combines precision and recall assessing the accuracy of a binary classification model.
3. SHAP Values[57]: SHAP values were used to explain the output of the machine learning model. This method employs a game-theoretic approach to quantify each feature’s contribution to the model’s predictions.

This comprehensive approach ensured a robust identification of the most significant features influencing the model’s performance.

## Declarations

### Consent for publication

#### Availability of data and materials

All data generated or analyzed during this study are included in this published article [and its supplementary information files].

#### Funding

This study was supported in part by a fellowship from the Edmond J. Safra Center for Bioinformatics at Tel-Aviv University, by the Debi Ofakim research fellowship by Miri and Efraim, and by Horizon 2020 ATTRACT project Re-sense. Research in the Belkin laboratory was partially supported by the Minerva Center for Bio-Hybrid Complex Systems.

## Supplementary

**Supplementary S1 – Table of all generated variants**

**Supplementary S2 – Table of the synthetic library data**

**Supplementary S3 – Table of all features names and description**

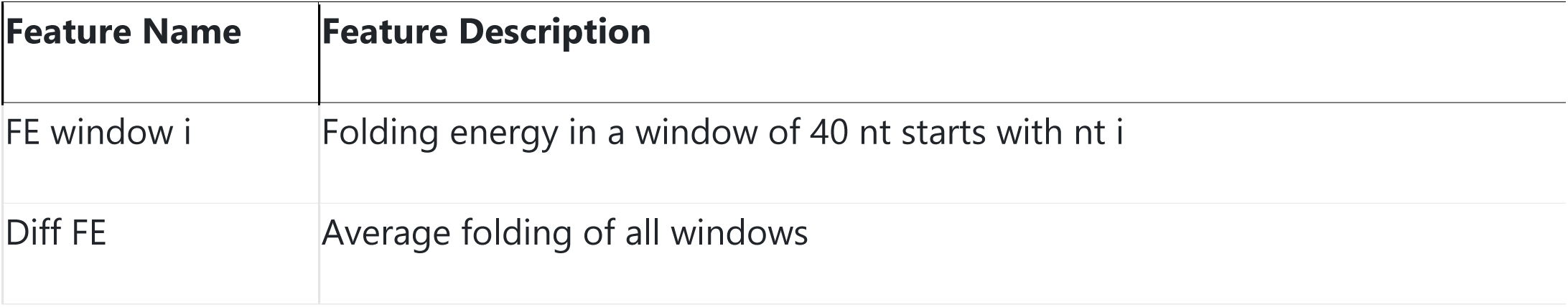
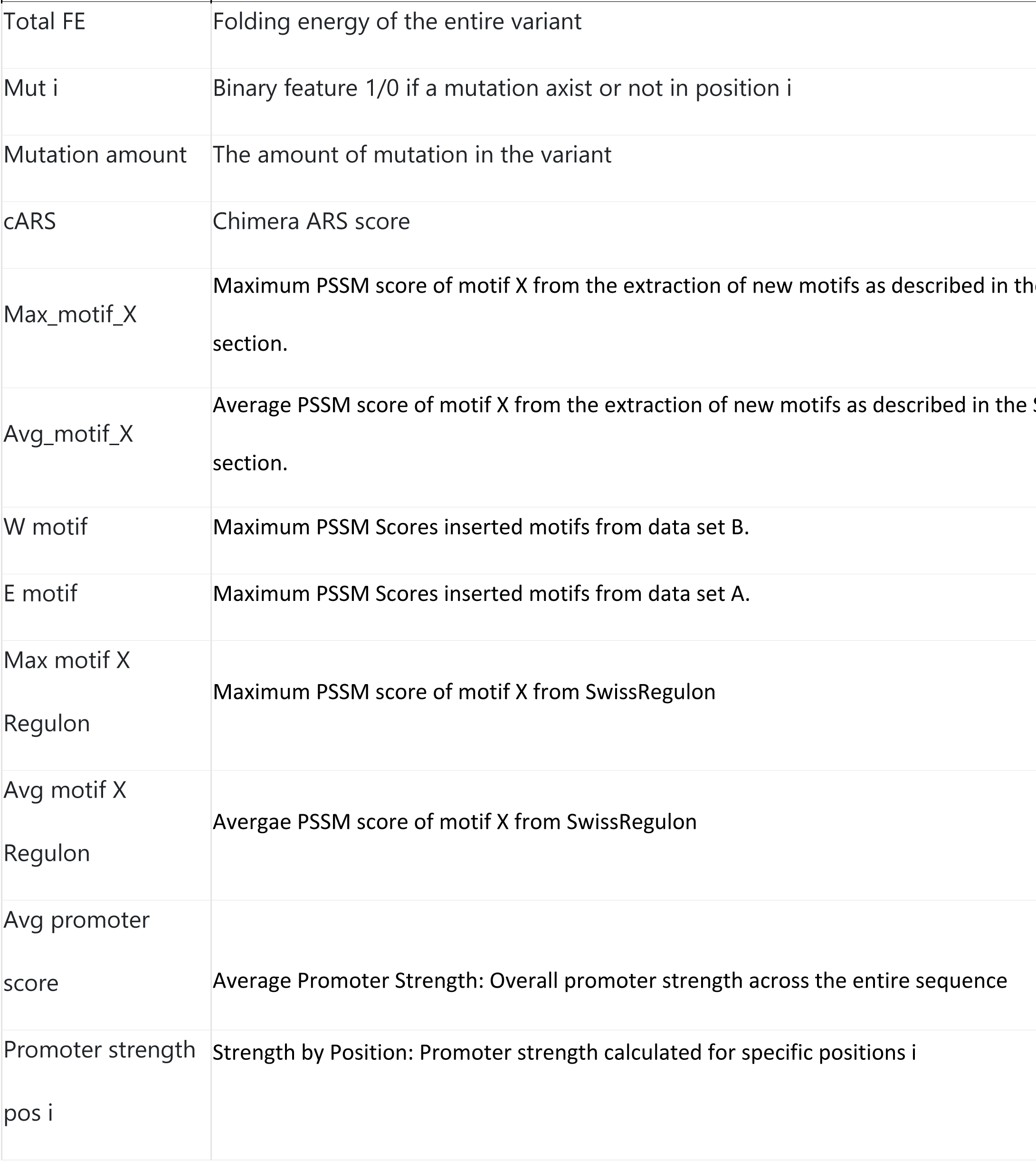
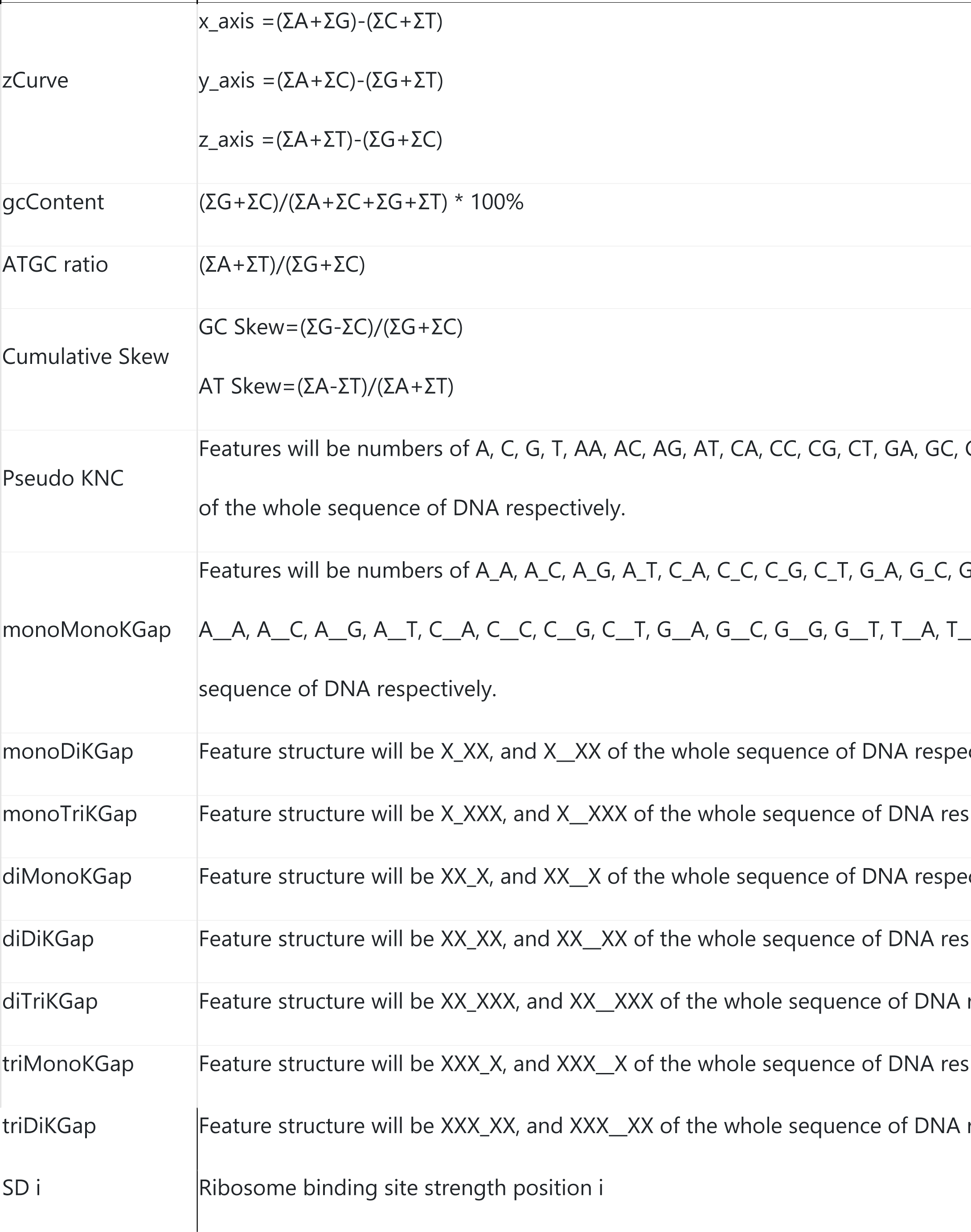

**Figure S1.**
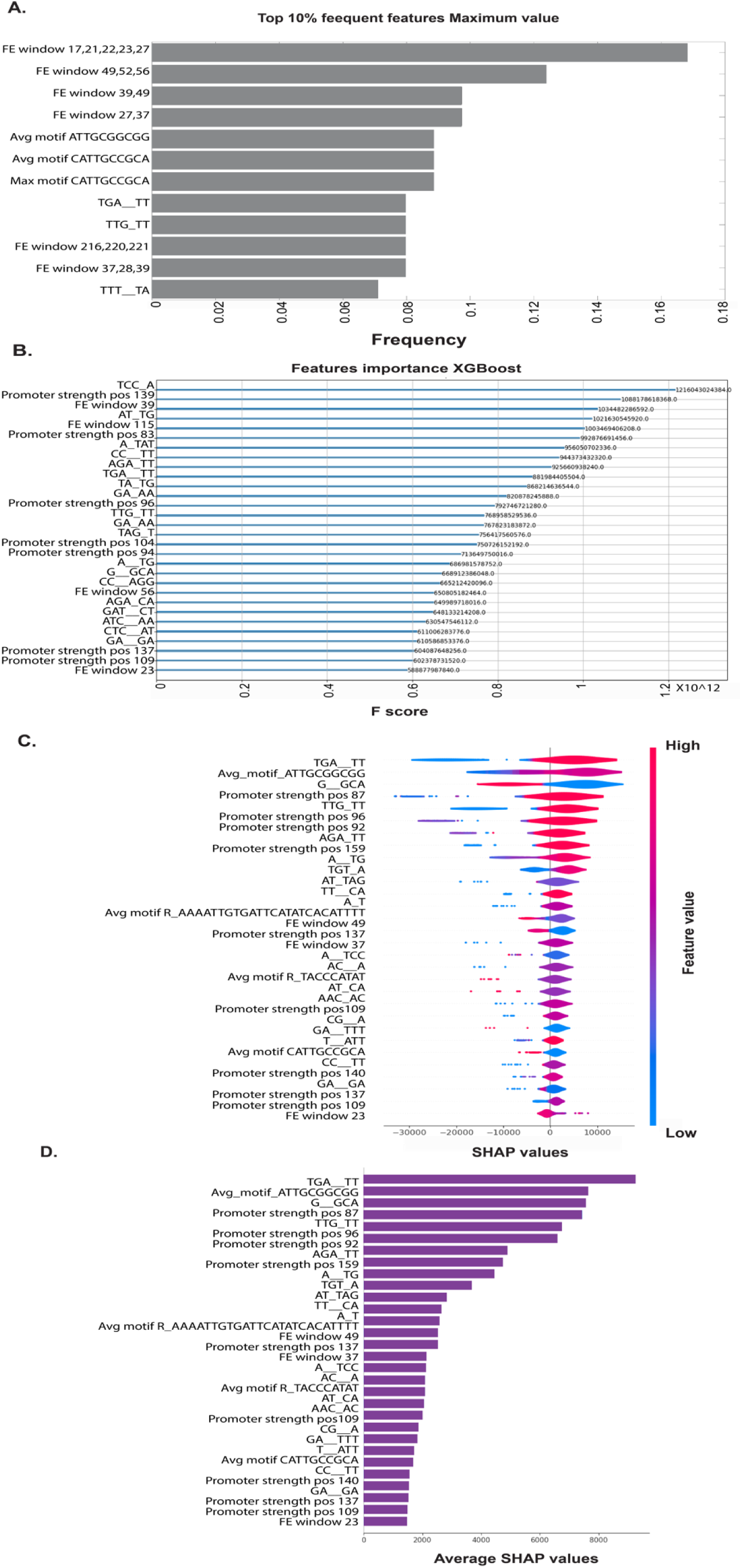
Most influential features predictor analysis Maximum value variable. **A**. Top 10% of frequent features from all cross-validation sets. **B.** Top 30 features F score. **C.** Top 30 features SHAP values and direction. D. Top 30 features averaged SHAP values.

**Figure S2.**
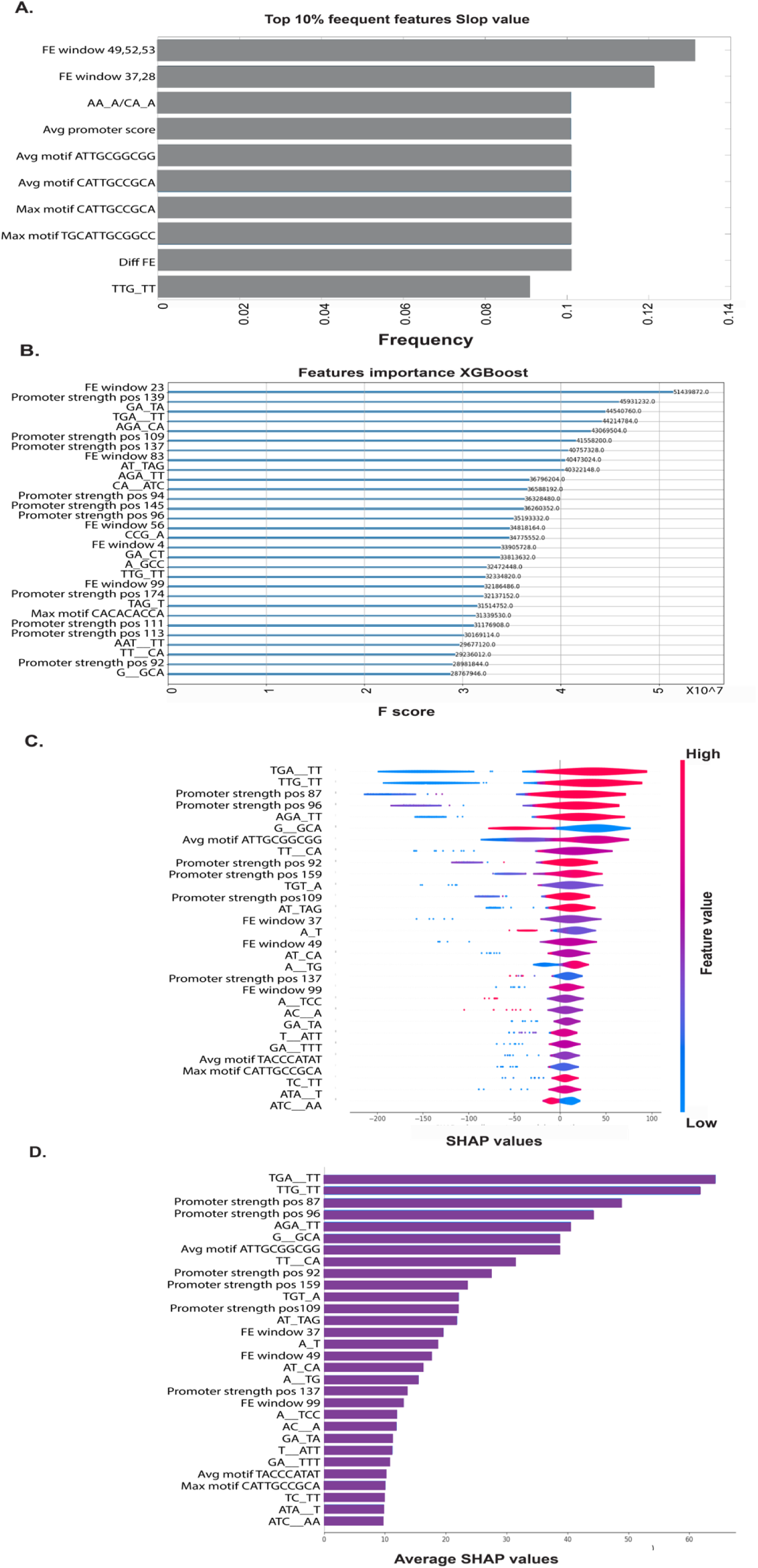
Most influential features predictor analysis Slop value variable. **A**. Top 10% of frequent features from all cross-validation sets. **B**. Top 30 features F score. **C**. Top 30 features SHAP values and direction. **D**. Top 30 features averaged SHAP values.

**Figure S3.**
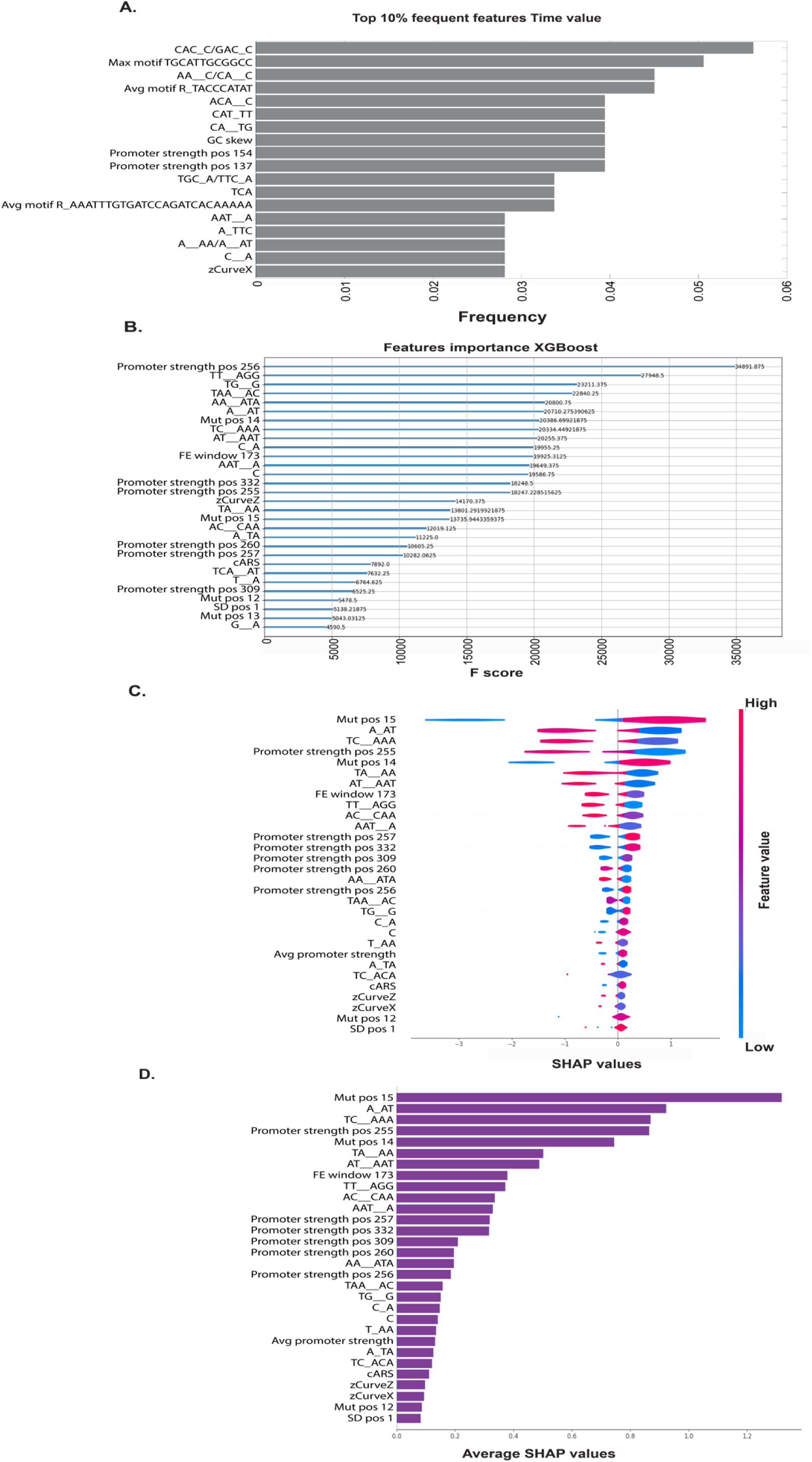
Most influential features predictor analysis Time to max value variable. **A**. Top 10% of frequent features from all cross-validation sets. **B**. Top 30 features F score. **C**. Top 30 features SHAP values and direction. **D**. Top 30 features averaged SHAP values.

**Figure S4.**
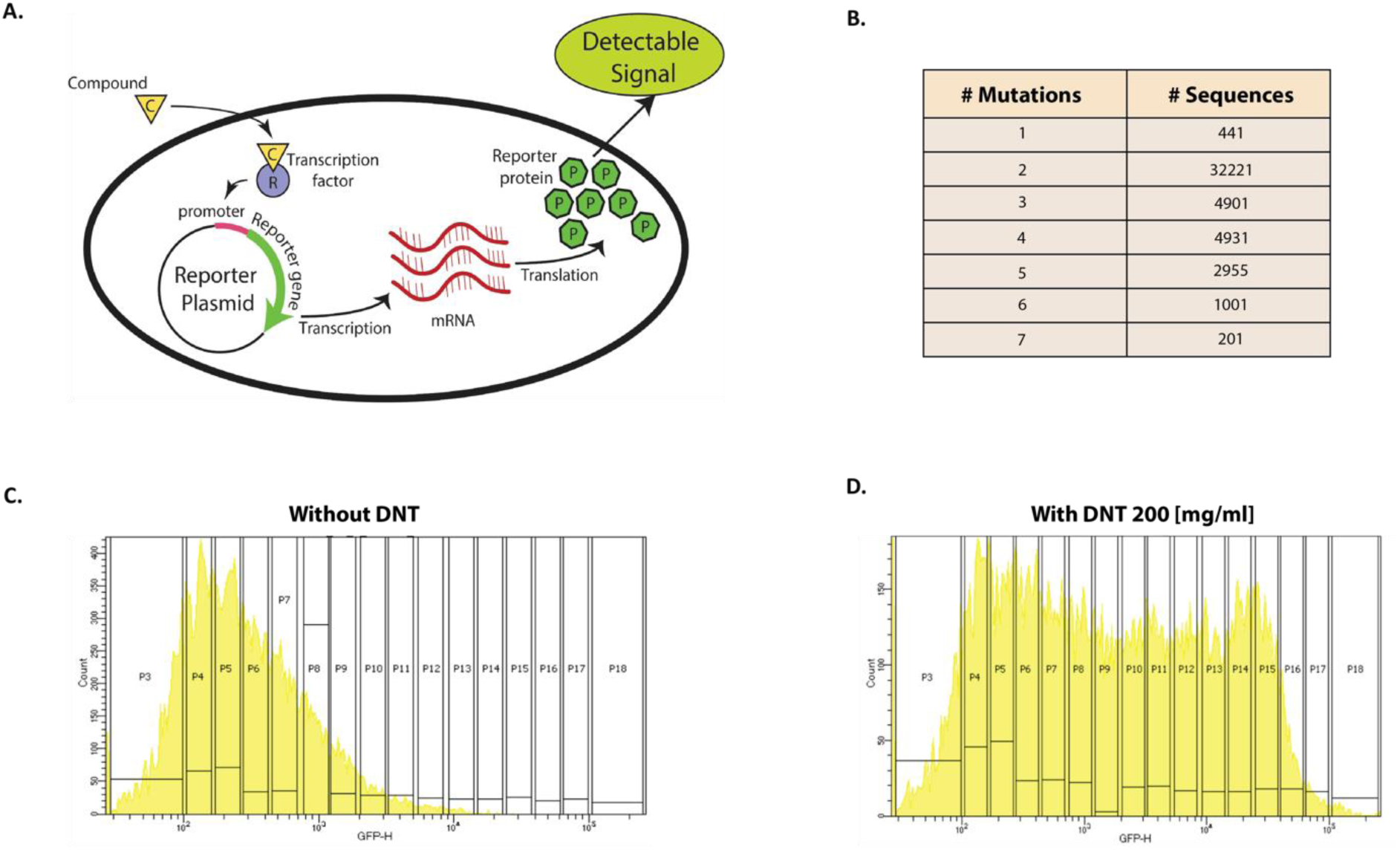
Synthetic library design. **A.** A promotor based biological sensor was the model system. The promotor was connected to a GFP (reporter gene) and the fluorescence levels was measured with and without exposure to DNT. **B.** The library was designed to contain all single mutations of, and a large sample of mutation pairs, triplets and higher order mutations as can be seen in the table. **C.** The library was sorted using FACS to 16 bins (in log scale) according to GFP fluorescence. The measured GFP fluoresces without DNT. **D.** The library was sorted using FACS to 16 bins (in log scale) according to GFP fluorescence. The measured GFP fluoresces in the presence of 200 μg/ml of DNT.

**Figure S5.**
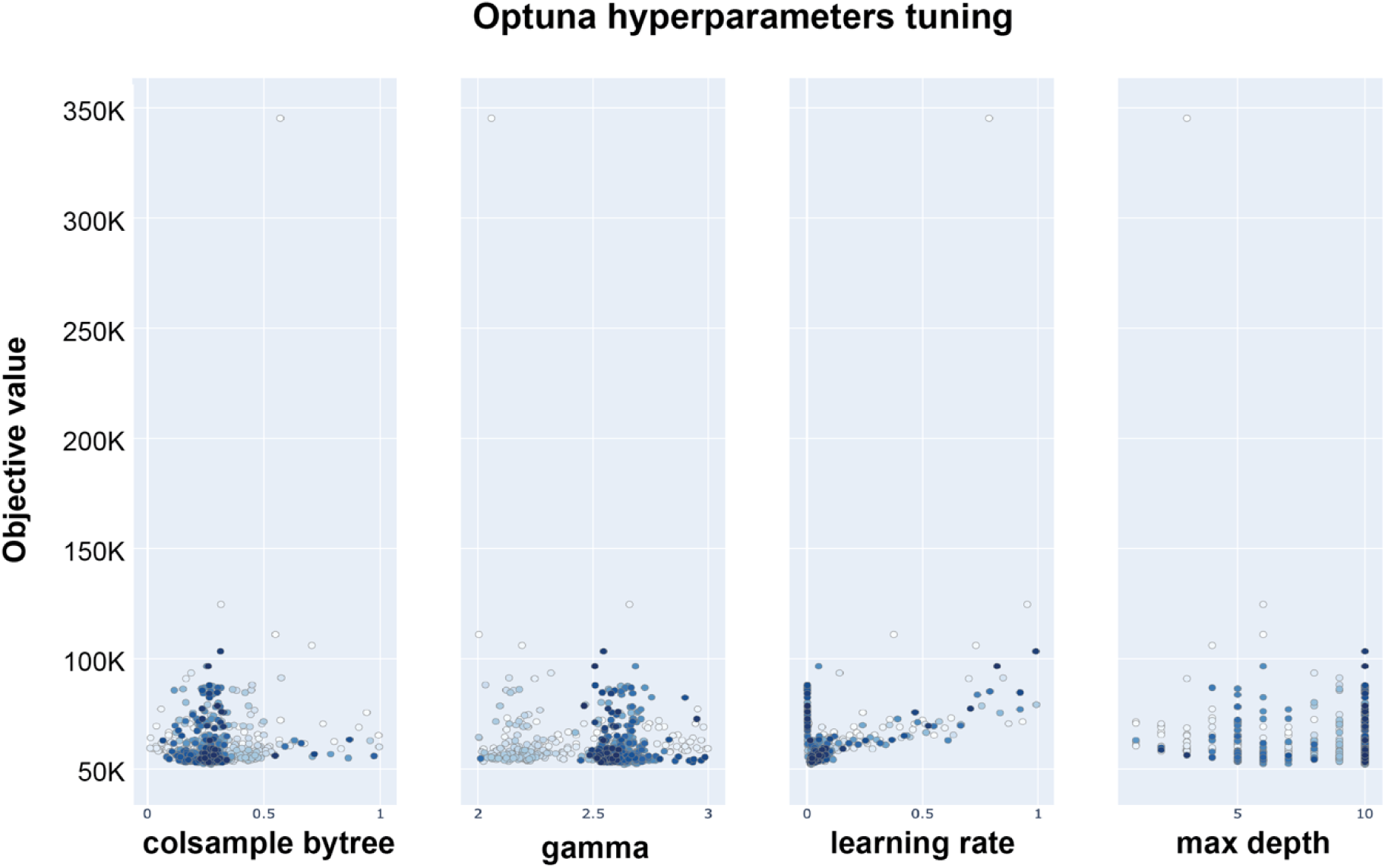
Hyperparameters optimization by Optuna. The optimization of hyperparameters using Optona involved systematically testing a range of values to identify the optimal settings for each parameter. As illustrated in the graph for a few parameters, Optona evaluated various hyperparameters, demonstrating the differences in performance across tested ranges. This process revealed that some hyperparameters had a broader range of values tested, which allowed us to assess their impact comprehensively. Based on these evaluations, it became evident that certain ranges could be narrowed down iteratively, focusing on the most promising intervals. By reducing the range and concentrating on these optimal intervals, we could more effectively pinpoint the best values for each hyperparameter, enhancing the overall model performance and efficiency.

