## Supplementary information for "Design of bacterial DNT sensors based on computational models"

**Supplementary S1 – Table of all generated variants**

**Supplementary S2 – Table of the synthetic library data**

**Supplementary S3 – Table of all features names and description**

| Feature Name | Feature Description |
|---|---|
| FE window i | Folding energy in a window of 40 nt starts with nt i |
| Diff FE | Average folding of all windows |
| Total FE | Folding energy of the entire variant |
| Mut i | Binary feature 1/0 if a mutation axist or not in position i |
| Mutation amount | The amount of mutation in the variant |
| cARS | Chimera ARS score |
| Max_motif_X | Maximum PSSM score of motif X from the extraction of new motifs as described i[illegible] Extraction section. |
| Avg_motif_X | Average PSSM score of motif X from the extraction of new motifs as described in [illegible] Extraction section. |
| W motif | Maximum PSSM Scores inserted motifs from data set B. |

| Feature Name | Feature Description |
|---|---|
| E motif | Maximum PSSM Scores inserted motifs from data set A. |
| Max motif X Regulon | Maximum PSSM score of motif X from SwissRegulon |
| Avg motif X Regulon | Avergae PSSM score of motif X from SwissRegulon |
| Avg promoter score | Average Promoter Strength: Overall promoter strength across the entire sequence |
| Promoter strength pos i | Strength by Position: Promoter strength calculated for specific positions i |
| zCurve | $\text{x\_axis} = (\Sigma A+\Sigma G)-(\Sigma C+\Sigma T)$<br>$\text{y\_axis} = (\Sigma A+\Sigma C)-(\Sigma G+\Sigma T)$<br>$\text{z\_axis} = (\Sigma A+\Sigma T)-(\Sigma G+\Sigma C)$ |
| gcContent | $(\Sigma G+\Sigma C)/(\Sigma A+\Sigma C+\Sigma G+\Sigma T) * 100\%$ |
| ATGC ratio | $(\Sigma A+\Sigma T)/(\Sigma G+\Sigma C)$ |
| Cumulative Skew | GC Skew$=(\Sigma G-\Sigma C)/(\Sigma G+\Sigma C)$<br>AT Skew$=(\Sigma A-\Sigma T)/(\Sigma A+\Sigma T)$ |
| Pseudo KNC | Features will be numbers of A, C, G, T, AA, AC, AG, AT, CA, CC, CG, CT, GA, GC, GG, [illegible] of the whole sequence of DNA respectively. |

| Feature Name | Feature Description |
|---|---|
| monoMonoKGap | Features will be numbers of A_A, A_C, A_G, A_T, C_A, C_C, C_G, C_T, G_A, G_C, G_G, A__A, A__C, A__G, A__T, C__A, C__C, C__G, C__T, G__A, G__C, G__G, G__T, T__A, T__C, sequence of DNA respectively. |
| monoDiKGap | Feature structure will be X_XX, and X__XX of the whole sequence of DNA respectiv |
| monoTriKGap | Feature structure will be X_XXX, and X__XXX of the whole sequence of DNA respec |
| diMonoKGap | Feature structure will be XX_X, and XX__X of the whole sequence of DNA respectiv |
| diDiKGap | Feature structure will be XX_XX, and XX__XX of the whole sequence of DNA respec |
| diTriKGap | Feature structure will be XX_XXX, and XX__XXX of the whole sequence of DNA resp |
| triMonoKGap | Feature structure will be XXX_X, and XXX__X of the whole sequence of DNA respec |
| triDiKGap | Feature structure will be XXX_XX, and XXX__XX of the whole sequence of DNA resp |
| SD i | Ribosome binding site strength position i |

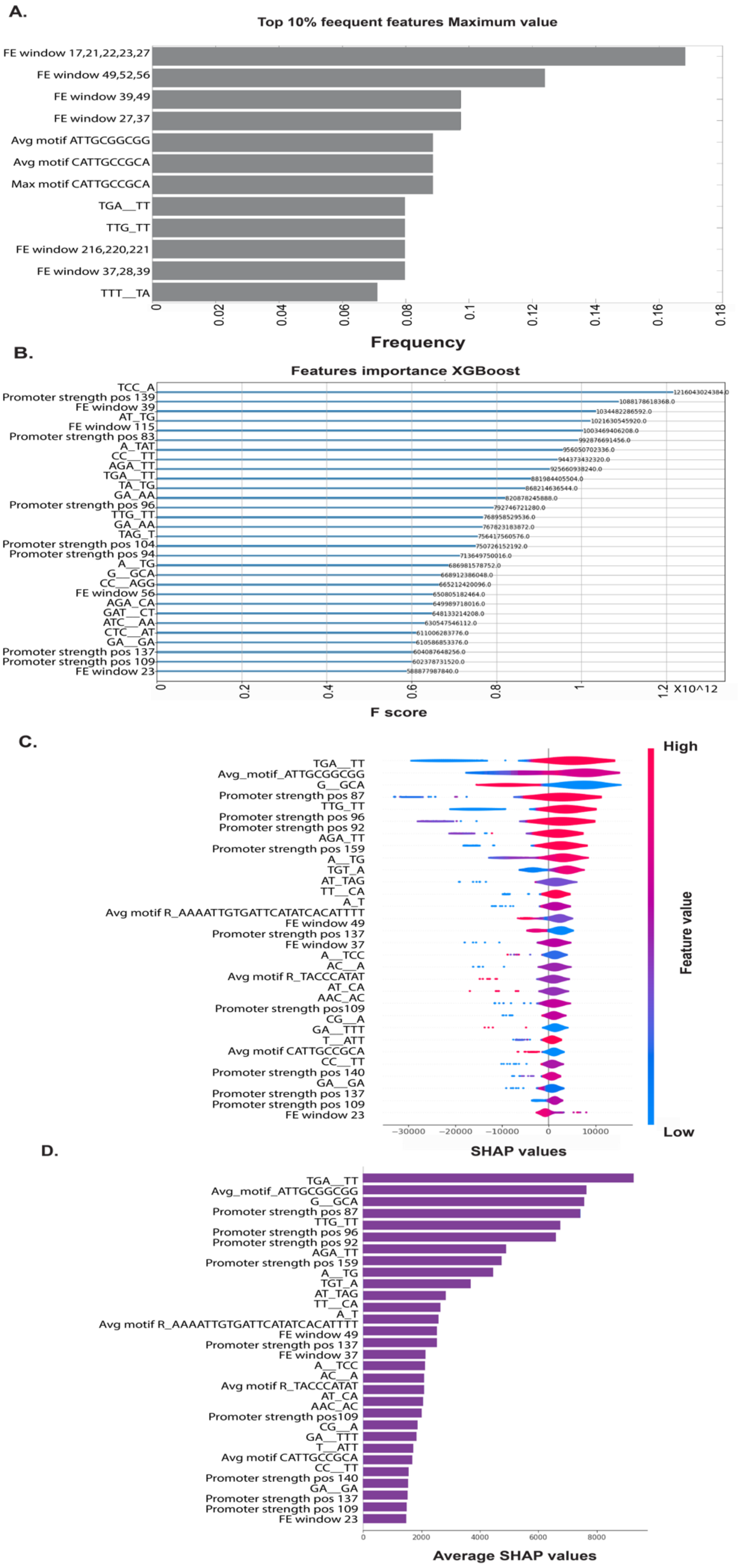
A.
Top 10% feequent features Maximum value
FE window 17,21,22,23,27
FE window 49,52,56
FE window 39,49
FE window 27,37
Avg motif ATTGCGGCGG
Avg motif CATTGCCGCA
Max motif CATTGCCGCA
TGA__TT
TTG_TT
FE window 216,220,221
FE window 37,28,39
TTT__TA
0 0.02 0.04 0.06 0.08 0.1 0.12 0.14 0.16 0.18
Frequency
B.
Features importance XGBoost
TCC_A 1216043024384.0
Promoter strength pos 139 1088178618368.0
FE window 39 1034482286592.0
AT_TG 1021630545920.0
FE window 115 1003469406208.0
Promoter strength pos 83 992876691456.0
A_TAT 956050702336.0
CC__TT 944373432320.0
AGA_TT 925660938240.0
TGA__TT 881984405504.0
TA_TG 868214636544.0
GA_AA 820878245888.0
Promoter strength pos 96 792746721280.0
TTG_TT 768958529536.0
GA_AA 767823183872.0
TAG_T 756417560576.0
Promoter strength pos 104 750726152192.0
Promoter strength pos 94 713649750016.0
A__TG 686981578752.0
G__GCA 668912386048.0
CC__AGG 665212420096.0
FE window 56 650805182464.0
AGA_CA 649989718016.0
GAT__CT 648133214208.0
ATC__AA 630547546112.0
CTC__AT 611006283776.0
GA__GA 610586853376.0
Promoter strength pos 137 604087648256.0
Promoter strength pos 109 602378731520.0
FE window 23 588877987840.0
0 0.2 0.4 0.6 0.8 1 1.2 X10^12
F score
C.
TGA__TT
Avg_motif_ATTGCGGCGG
G__GCA
Promoter strength pos 87
TTG_TT
Promoter strength pos 96
Promoter strength pos 92
AGA_TT
Promoter strength pos 159
A__TG
TGT_A
AT_TAG
TT__CA
A_T
Avg motif R_AAAATTGTGATTCATATCACATTTT
FE window 49
Promoter strength pos 137
FE window 37
A__TCC
AC__A
Avg motif R_TACCCATAT
AT_CA
AAC_AC
Promoter strength pos109
CG__A
GA__TTT
T__ATT
Avg motif CATTGCCGCA
CC__TT
Promoter strength pos 140
GA__GA
Promoter strength pos 137
Promoter strength pos 109
FE window 23
−30000 −20000 −10000 0 10000
SHAP values
High
Feature value
Low
D.
TGA__TT
Avg_motif_ATTGCGGCGG
G__GCA
Promoter strength pos 87
TTG_TT
Promoter strength pos 96
Promoter strength pos 92
AGA_TT
Promoter strength pos 159
A__TG
TGT_A
AT_TAG
TT__CA
A_T
Avg motif R_AAAATTGTGATTCATATCACATTTT
FE window 49
Promoter strength pos 137
FE window 37
A__TCC
AC__A
Avg motif R_TACCCATAT
AT_CA
AAC_AC
Promoter strength pos109
CG__A
GA__TTT
T__ATT
Avg motif CATTGCCGCA
CC__TT
Promoter strength pos 140
GA__GA
Promoter strength pos 137
Promoter strength pos 109
FE window 23
0 2000 4000 6000 8000
Average SHAP values

***Figure S1.*** *Most influential features predictor analysis Maximum value variable*. ***A.*** *Top 10% of frequent features from all cross-validation sets.* ***B.*** *Top 30 features F score.* ***C.*** *Top 30 features SHAP values and direction.* ***D.*** *Top 30 features averaged SHAP values.*

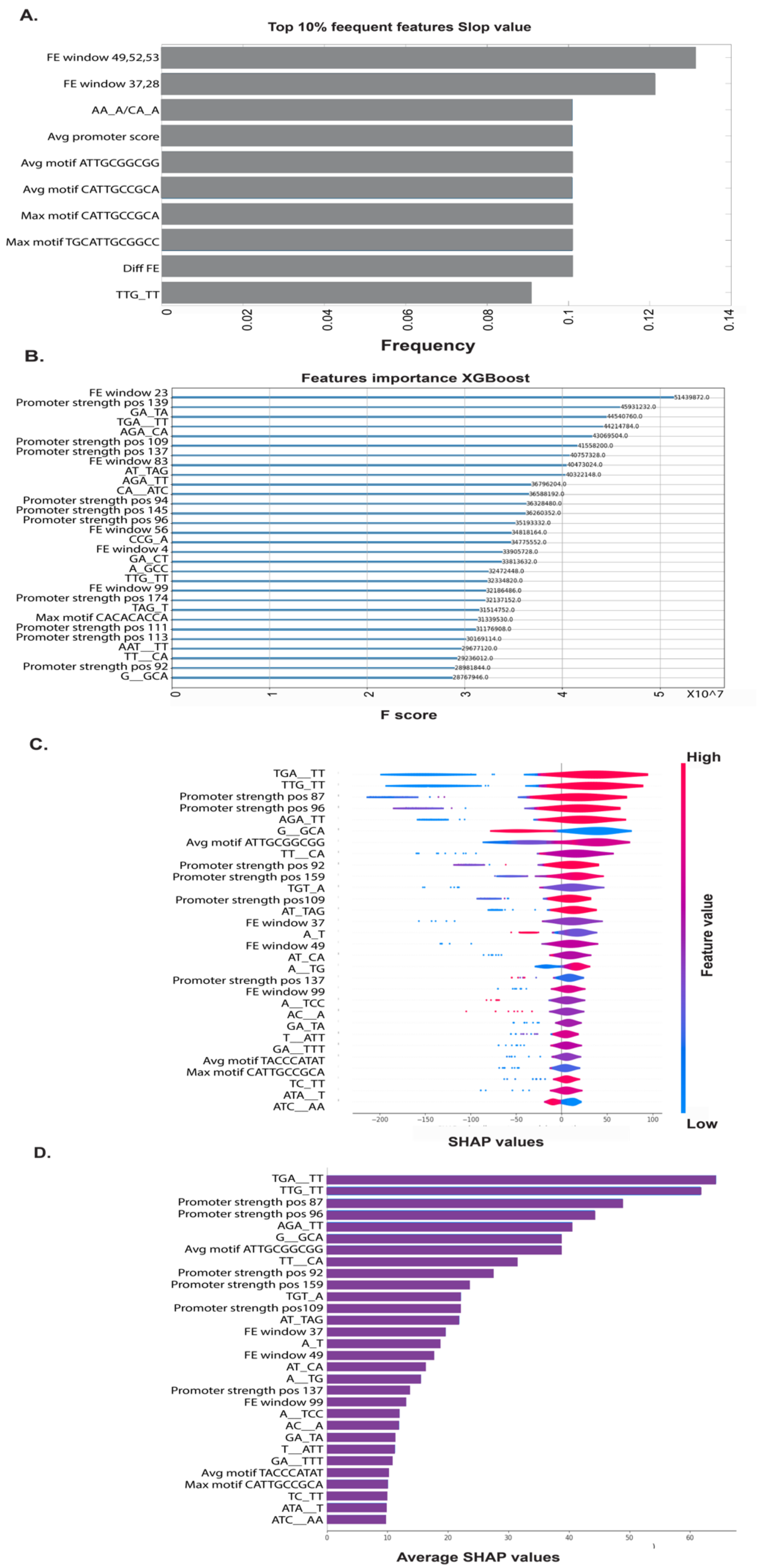
A.
Top 10% feequent features Slop value
FE window 49,52,53
FE window 37,28
AA_A/CA_A
Avg promoter score
Avg motif ATTGCGGCGG
Avg motif CATTGCCGCA
Max motif CATTGCCGCA
Max motif TGCATTGCGGCC
Diff FE
TTG_TT
0
0.02
0.04
0.06
0.08
0.1
0.12
0.14
Frequency
B.
Features importance XGBoost
FE window 23
Promoter strength pos 139
GA_TA
TGA__TT
AGA_CA
Promoter strength pos 109
Promoter strength pos 137
FE window 83
AT_TAG
AGA_TT
CA__ATC
Promoter strength pos 94
Promoter strength pos 145
Promoter strength pos 96
FE window 56
CCG_A
FE window 4
GA_CT
A_GCC
TTG_TT
FE window 99
Promoter strength pos 174
TAG_T
Max motif CACACACCA
Promoter strength pos 111
Promoter strength pos 113
AAT__TT
TT__CA
Promoter strength pos 92
G__GCA
51439872.0
45931232.0
44540760.0
44214784.0
43069504.0
41558200.0
40757328.0
40473024.0
40322148.0
36796204.0
36588192.0
36328480.0
36260352.0
35193332.0
34818164.0
34775552.0
33905728.0
33813632.0
32472448.0
32334820.0
32186486.0
32137152.0
31514752.0
31339530.0
31176908.0
30169114.0
29677120.0
29236012.0
28981844.0
28767946.0
0
1
2
3
4
5
X10^7
F score
C.
TGA__TT
TTG_TT
Promoter strength pos 87
Promoter strength pos 96
AGA_TT
G__GCA
Avg motif ATTGCGGCGG
TT__CA
Promoter strength pos 92
Promoter strength pos 159
TGT_A
Promoter strength pos109
AT_TAG
FE window 37
A_T
FE window 49
AT_CA
A__TG
Promoter strength pos 137
FE window 99
A__TCC
AC__A
GA_TA
T__ATT
GA__TTT
Avg motif TACCCATAT
Max motif CATTGCCGCA
TC_TT
ATA__T
ATC__AA
−200
−150
−100
−50
0
50
100
SHAP values
High
Feature value
Low
D.
TGA__TT
TTG_TT
Promoter strength pos 87
Promoter strength pos 96
AGA_TT
G__GCA
Avg motif ATTGCGGCGG
TT__CA
Promoter strength pos 92
Promoter strength pos 159
TGT_A
Promoter strength pos109
AT_TAG
FE window 37
A_T
FE window 49
AT_CA
A__TG
Promoter strength pos 137
FE window 99
A__TCC
AC__A
GA_TA
T__ATT
GA__TTT
Avg motif TACCCATAT
Max motif CATTGCCGCA
TC_TT
ATA__T
ATC__AA
0
10
20
30
40
50
60
Average SHAP values

***Figure S2.*** *Most influential features predictor analysis Slop value variable*. ***A.*** *Top 10% of frequent features from all cross-validation sets.* ***B.*** *Top 30 features F score.* ***C.*** *Top 30 features SHAP values and direction.* ***D.*** *Top 30 features averaged SHAP values.*

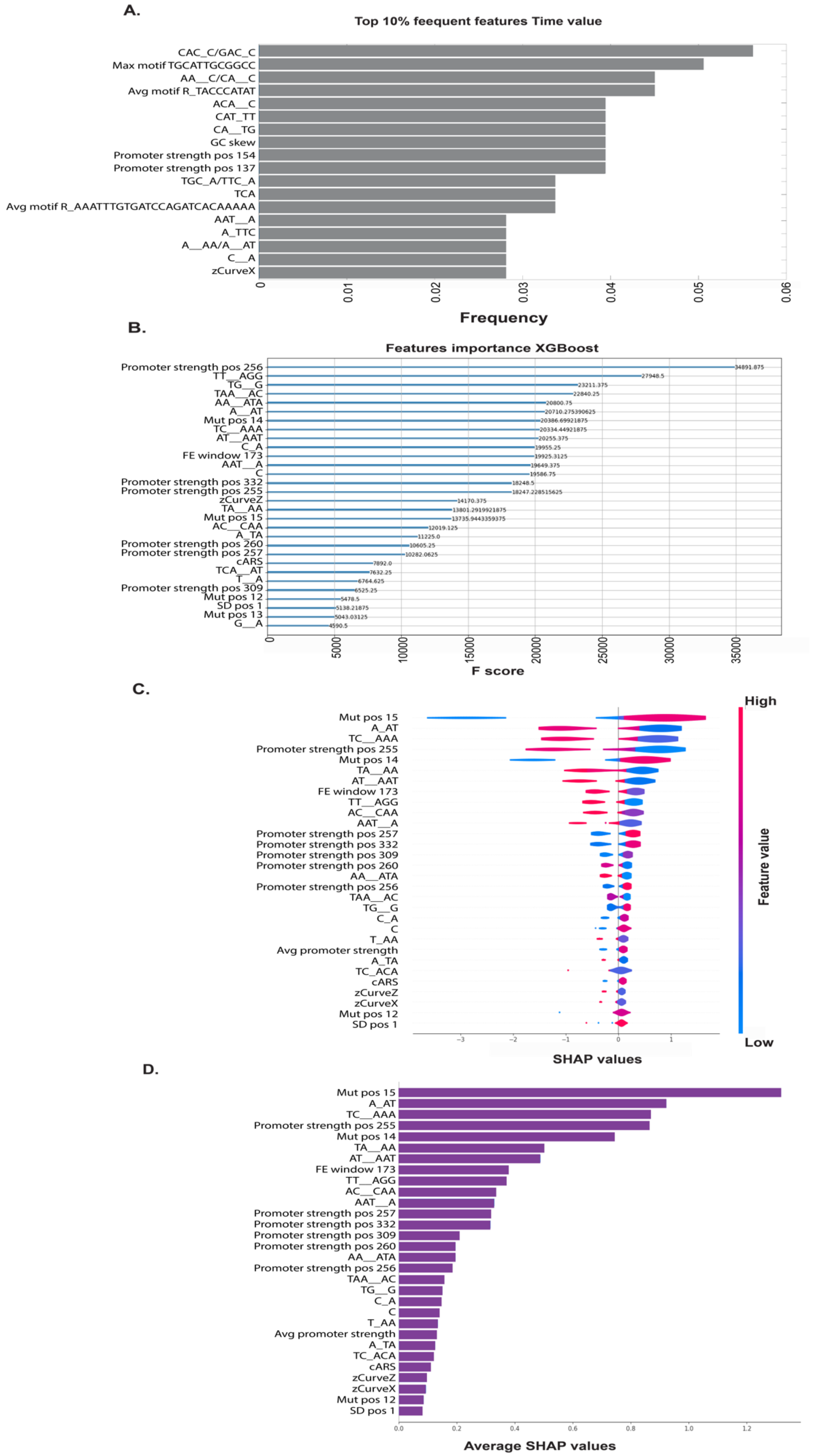

A.
Top 10% feequent features Time value
CAC_C/GAC_C
Max motif TGCATTGCGGCC
AA__C/CA__C
Avg motif R_TACCCATAT
ACA__C
CAT_TT
CA__TG
GC skew
Promoter strength pos 154
Promoter strength pos 137
TGC_A/TTC_A
TCA
Avg motif R_AAATTTGTGATCCAGATCACAAAAA
AAT__A
A_TTC
A__AA/A__AT
C__A
zCurveX
0
0.01
0.02
0.03
0.04
0.05
0.06
Frequency
B.
Features importance XGBoost
Promoter strength pos 256
TT__AGG
TG__G
TAA__AC
AA__ATA
A__AT
Mut pos 14
TC__AAA
AT__AAT
C_A
FE window 173
AAT__A
C
Promoter strength pos 332
Promoter strength pos 255
zCurveZ
TA__AA
Mut pos 15
AC__CAA
A_TA
Promoter strength pos 260
Promoter strength pos 257
cARS
TCA__AT
T__A
Promoter strength pos 309
Mut pos 12
SD pos 1
Mut pos 13
G__A
34891.875
27948.5
23211.375
22840.25
20800.75
20710.275390625
20386.69921875
20334.44921875
20255.375
19955.25
19925.3125
19649.375
19586.75
18248.5
18247.228515625
14170.375
13801.2919921875
13735.9443359375
12019.125
11225.0
10605.25
10282.0625
7892.0
7632.25
6764.625
6525.25
5478.5
5138.21875
5043.03125
4590.5
0
5000
10000
15000
20000
25000
30000
35000
F score
C.
Mut pos 15
A_AT
TC__AAA
Promoter strength pos 255
Mut pos 14
TA__AA
AT__AAT
FE window 173
TT__AGG
AC__CAA
AAT__A
Promoter strength pos 257
Promoter strength pos 332
Promoter strength pos 309
Promoter strength pos 260
AA__ATA
Promoter strength pos 256
TAA__AC
TG__G
C_A
C
T_AA
Avg promoter strength
A_TA
TC_ACA
cARS
zCurveZ
zCurveX
Mut pos 12
SD pos 1
−3
−2
−1
0
1
SHAP values
High
Feature value
Low
D.
Mut pos 15
A_AT
TC__AAA
Promoter strength pos 255
Mut pos 14
TA__AA
AT__AAT
FE window 173
TT__AGG
AC__CAA
AAT__A
Promoter strength pos 257
Promoter strength pos 332
Promoter strength pos 309
Promoter strength pos 260
AA__ATA
Promoter strength pos 256
TAA__AC
TG__G
C_A
C
T_AA
Avg promoter strength
A_TA
TC_ACA
cARS
zCurveZ
zCurveX
Mut pos 12
SD pos 1
0.0
0.2
0.4
0.6
0.8
1.0
1.2
Average SHAP values

***Figure S3.*** *Most influential features predictor analysis Time to max value variable*. ***A.*** *Top 10% of frequent features from all cross-validation sets.* ***B.*** *Top 30 features F score.* ***C.*** *Top 30 features SHAP values and direction.* ***D.*** *Top 30 features averaged SHAP values.*

A.

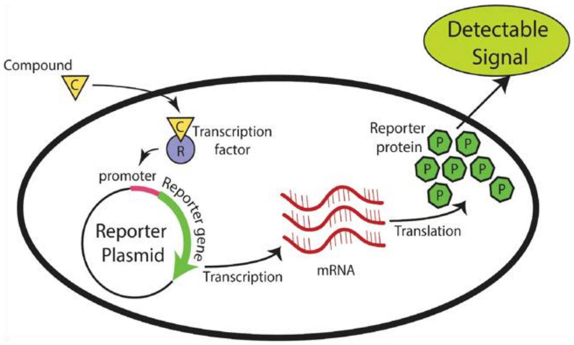


B.

| # Mutations | # Sequences |
|---|---|
| 1 | 441 |
| 2 | 32221 |
| 3 | 4901 |
| 4 | 4931 |
| 5 | 2955 |
| 6 | 1001 |
| 7 | 201 |

C.

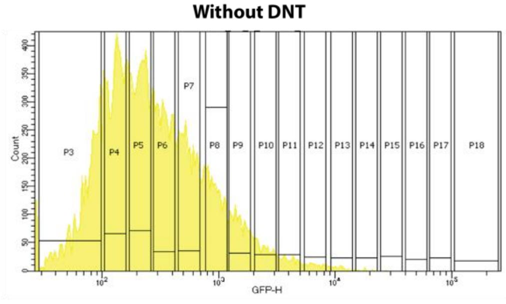


D.

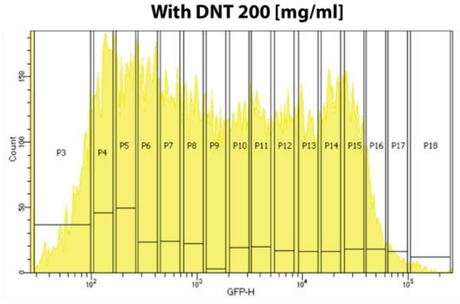


***Figure S4. Synthetic library design. A.*** *A promotor-based biological sensor was the model system. The promotor was connected to a GFP (reporter gene) and the fluorescence levels were measured with and without exposure to DNT.* ***B.*** *The library was designed to contain all single mutations, and a large sample of mutation pairs, triplets, and higher-order mutations can be seen in the table.* ***C.*** *The library was sorted using FACS into 16 bins (on a log scale) according to GFP fluorescence. The measured GFP*

*fluoresces without DNT.* ***D.*** *The library was sorted using FACS into 16 bins (in log scale) according to GFP fluorescence. The measured GFP fluoresces in the presence of 200 μg/ml of DNT.*

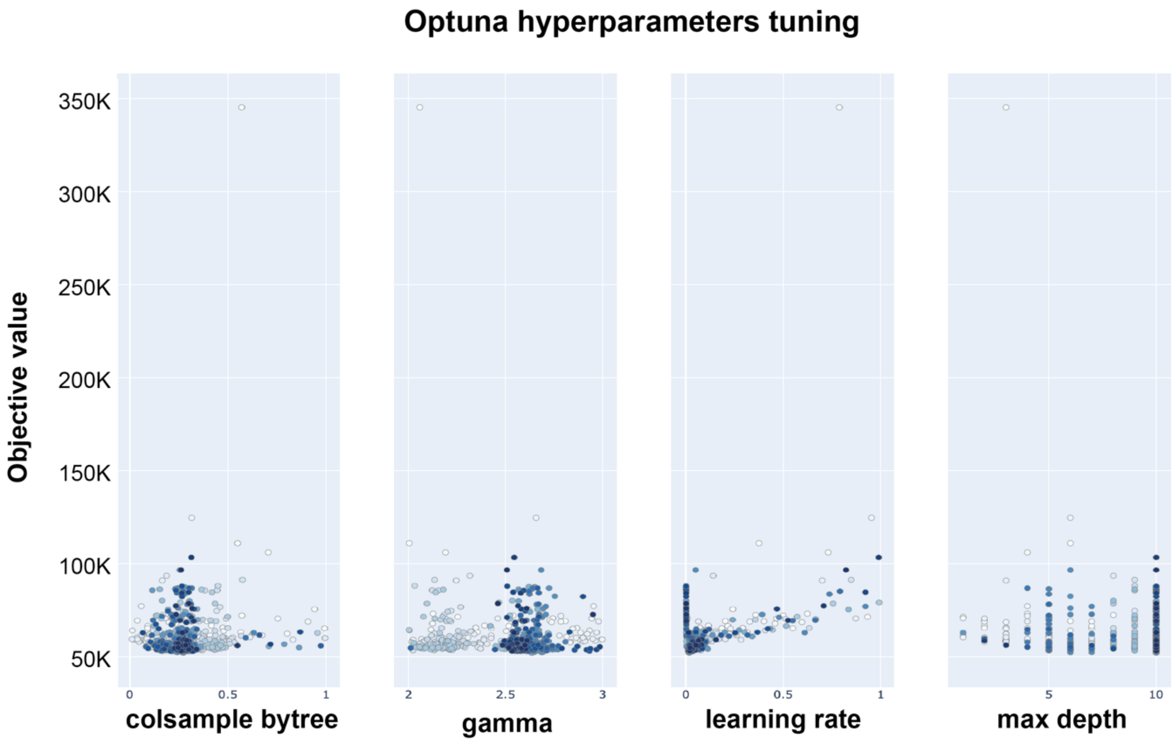


***Figure S5. Hyperparameters optimization by Optuna.*** *The optimization of hyperparameters using Optona involved systematically testing a range of values to identify the optimal settings for each parameter. As illustrated in the graph for a few parameters, Optona evaluated various hyperparameters, demonstrating the differences in performance across tested ranges. This process revealed that some hyperparameters had a broader range of values tested, which allowed us to assess their impact comprehensively. Based on these evaluations, it became evident that certain ranges*

*could be narrowed down iteratively, focusing on the most promising intervals. By reducing the range and concentrating on these optimal intervals, we could more effectively pinpoint the best values for each hyperparameter, enhancing the overall model performance and efficiency.*
